# The effects of weak selection on neutral diversity at linked sites

**DOI:** 10.1101/2021.11.27.470208

**Authors:** Brian Charlesworth

## Abstract

The effects of selection on variability at linked sites have an important influence on levels and patterns of within-population variation across the genome. Most theoretical models of these effects have assumed that selection is sufficiently strong that allele frequency changes at the loci concerned are largely deterministic. These models have led to the conclusion that directional selection for selectively favorable mutations, or against recurrent deleterious mutations, reduces nucleotide site diversity at linked neutral sites. Recent work has shown, however, that fixations of weakly selected mutations, accompanied by significant stochastic changes in allele frequencies, can sometimes cause higher diversity at linked sites when compared with the effects of fixations of neutral mutations. The present paper extends this work by deriving approximate expressions for the mean conditional times to fixation and loss of mutations subject to selection, and analysing the conditions under which selection increases rather than reduces these times. Simulations are used to examine the relations between diversity at a neutral site and the fixation and loss times of mutations at a linked site that is subject to selection. It is shown that the long-term level of neutral diversity can be increased over the purely neutral value by recurrent fixations and losses of linked, weakly selected dominant or partially dominant favorable mutations, or linked recessive or partially recessive deleterious mutations. The results are used to examine the conditions under which associative overdominance, as opposed to background selection, is likely to operate.

## Introduction

There is now a large body of data showing that the levels of within-population DNA sequence diversity across the genomes of many organisms are significantly affected by the effects of selection on sites linked to those under observation, especially in genomic regions or species where recombination rates are low; for recent reviews, see Cutter and Payseur (2013) and Charlesworth and Jensen (2021). Interpretations of these observations have mainly focussed on reductions in variation at linked neutral or nearly neutral sites caused by the spread of selectively favorable variants (selective sweeps), or the elimination of rare, deleterious mutations (background selection). The population genetic models used to describe these processes usually assume that selection is sufficiently strong in relation to genetic drift that deterministic equations are sufficient to describe the behavior of the sites under selection, except for the initial and final periods of the fixation of beneficial mutations (reviewed by Charlesworth and Jensen 2021).

Interest has, however, recently been revived in the process known as associative overdominance (Frydenberg 1963; Sved 1968; Ohta 1971; Latter 1998; Pamilo and Palsson 1998, 1999; Wang and Hill 1999; Zhao and Charlesworth 2016; Waller 2021), whereby the level of diversity at a neutral locus in a diploid population can be enhanced by the presence of linked deleterious alleles maintained by mutation pressure. Recent theoretical work has shown that this can happen when the deleterious alleles concerned are sufficiently recessive, and selection is sufficiently weak in relation to drift (Zhao and Charlesworth 2016). The latter condition requires the product of the effective size of the population (*N*_*e*_) and the selection coefficient against homozygotes for a mutation (*s*) to be the order of 1 or less, consistent with previous results from computer simulations (Latter 1998; Palsson and Pamilo 1999; Wang and Hill 1999). There is evidence for the operation of associative overdominance (AOD) in both small populations (Latter 1998; Zhao and Charlesworth 2016; Schou et al. 2017; Waller 2021), and in genomic regions with low recombination, where *N*_*e*_ is reduced as a result of background selection and selective sweeps (Becher et al. 2020; Gilbert et al. 2020). In contrast, if the effects of drift on the frequencies of deleterious alleles are negligible, background selection (BGS) operates, causing a reduction in variability at linked sites (Charlesworth et al. 1993; Hudson and Kaplan 1995; Nordborg et al. 1996).

Under conditions where AOD is acting, deleterious variants at sites under selection are likely to become fixed as a result of drift, and reverse mutations that increase fitness can eventually arise and replace them as a result of the joint effects of drift and selection. In the long term, a population that is constant in size will reach a stochastic equilibrium under the joint effects of drift, mutation and selection, a situation that is similar to that envisaged in the Li-Bulmer model of the evolution of codon usage bias, which assumes a constant flux of fixations of favorable and beneficial mutations at sites under selection (Li 1987; Bulmer 1991; McVean and Charlesworth 1999). In order to understand the effects of AOD, it is therefore important to have a model of the effects of this flux on variability at linked neutral sites.

A basis for such a model is provided by the results of Mafessoni and Lachmann (2015) on the expected times to fixation and loss of new autosomal mutations in a randomly mating population, and on the effects of fixation events on the patterns of variability at linked neutral loci. They showed that a weakly selected (*N*_*e*_*s* of order 1), favorable mutation destined for fixation can have a longer mean time to fixation or loss than a neutral mutation, provided that it is dominant or partially dominant. The same applies to weakly selected, partially recessive deleterious mutations. Furthermore, the fixation of a dominant or partially dominant favorable mutation, or of a recessive or partially recessive deleterious mutation, can enhance variability at a linked neutral site compared with the effect of fixation of a neutral variant, although variability is still lower than in an equilibrium population without any selection. These conclusions have been confirmed in the simulations described by Johri *et al*. (2021); similar situations where fitnesses fluctuate over time have been studied by Kaushik and Jain (2021). As pointed out by Charlesworth and Jensen (2021), these results are relevant to the analysis of AOD by Zhao and Charlesworth (2016), who showed that a neutral locus closely linked to a locus subject to mutation to deleterious alleles can lose variability more slowly than in the absence of selection, provided that the mutations are recessive or partially recessive and selection is sufficiently weak relative to the effects of genetic drift.

The purpose of this paper is to obtain approximate analytical expressions for the expected sojourn times of new mutations for the case of weak selection, and to use these to illuminate the results of Mafessoni and Lachmann (2015) on the fixation and loss of weakly selected mutations. Computer simulations are used in conjunction with these results to examine the effects of fixations and losses of weakly selected mutations on variability at linked neutral sites. The results are used to develop a semi-analytical model of AOD for the case of stochastic equilibrium between mutation, selection and drift. They also provide a new way of describing BGS, when applied to situations when selection is so strong in relation to drift that deleterious mutations have a negligible chance of becoming fixed in the population. The focus is on the case when there is no recombination between neutral and selected sites, since this gives the clearest signal and allows the causes of the observed patterns to be analysed without the complications introduced by recombination.

## Material and methods

### Simulation methods

To check the accuracy of the diffusion equation results for expected times to fixation or loss (see the section *Theoretical Results*), a biallelic autosomal locus in a Wright-Fisher population with constant size *N* was modeled, implying that the effective population size *N*_*e*_ is equal to *N*. For a given simulation run, a single A_2_ allele was introduced into the population that was fixed for its alternative allele A_1_. The mutant allele could be either selectively favorable or deleterious. The expected change in the frequency *x* of A_2_ in a given generation for an assigned selection model was calculated using the standard discrete-generation selection formulation (for details of the model of selection, see the section *Theoretical Results, Approximate times to fixation and loss of a new mutation*). Binomial sampling using the frequency of A_2_ after selection and 2*N* as parameters was used to obtain the value of *x* in the next generation. Large numbers of replicate simulations were run in order to obtain the mean times to fixation and loss of A_2_, conditioned on its fixation or loss, respectively.

This procedure was modified in order to calculate the effects of a sweep on pairwise diversity at a neutral locus with an arbitrary degree of linkage to the selected locus. As described in Charlesworth (2020a) and Johri et al. (2021), the algorithm of Tajima (1990) was used to calculate the effects of a sweep on pairwise diversity at a neutral locus with an arbitrary degree of linkage to a selected locus with two alleles, A_1_ and A_2_. Tajima’s Equations (27) provide three coupled, forward-in-time recurrence relations for the expected diversities at the neutral locus for pairs of haplotypes carrying either A_1_ or A_2_, and the divergence between A_1_ and A_2_ haplotypes. These are conditioned on a given generation-by-generation trajectory of allele frequencies at the selected locus, assuming the infinite sites model of mutation and drift (Kimura 1971). In the present study, there is interest in the effects of both losses and fixations of either deleterious or advantageous A_2_ mutations on diversity statistics over the time-course leading to loss or fixation of A_2_, whereas the previous studies only considered the effects of fixations.

For a given simulation run, a single A_2_ allele was introduced into the population, with zero expected pairwise diversity at the associated neutral locus. The initial expected pairwise diversity among A_1_ alleles and divergence between A_1_ and A_2_ were set equal to those for an equilibrium population in the absence of selection, *θ* = 4*N*_*e*_*µ*, where *µ* is the neutral mutation rate. Since only diversities relative to *θ* are of interest here, *θ* was set to 0.001 in order to satisfy the infinite sites assumption. Equations (27) of Tajima (1990) were applied to the previous value of the allele frequency in order to obtain the state of the neutral locus in the new generation.

The simulation procedure for selection and drift at the selected locus was repeated generation by generation until A_2_ was lost or fixed; if the effect of loss on diversity at the neutral locus was of interest, only runs in which A_2_ was lost were retained, and the value of the pairwise diversity among A_2_ alleles at the time of its loss was determined. Similarly, the effects of fixations were studied by recording the properties of runs in which A_2_ was fixed. Large numbers of replicate simulations were used (between 10^4^ and 10^6^, depending on the parameter values), because there is a large amount of variation in the values of the population statistics between replicate runs.

Diversity statistics were obtained in each generation for the three genotypes at the selected locus. They were measured relative to the equilibrium neutral values, and were thus equivalent to the mean coalescent times on the timescale of 2*N*_*e*_ generations. The sums of the pairwise diversity statistics were taken over the whole time course of a loss or fixation, for A_1_ versus A_1_, A_1_ versus A_2_, and A_2_ versus A_2_ haplotypes, as well as the mean over all three of these comparisons, weighted by their frequencies in the generation in question. In addition, the neutral diversity of fixed haplotypes at the time of loss or fixation of the mutation was determined.

### Numerical integrations

The notation and results described in Ewens (2004, Chapters 4 and 5) are used here, with some slight modifications. The diffusion equation expressions shown in the Appendix yield the sojourn time densities at frequency *x* of A_2_ given an initial frequency *q*, which are denoted by *t*\*(*x, q*) and *t*\*\*(*x, q*) for fixations and losses conditioned on fixation or loss, respectively (see Equations A1 and A6). The mean times spent between frequencies *x* and *x* + d*x* are given by *t*\*(*x, q*) d*x* and *t*\*\*(*x, q*) d*x*, respectively, where d*x* is an arbitrarily small increment in *x*. The sojourn time densities involve integrals of the function*ψ*(*y*), defined by Equation (A1c). For a given value of *x*, these integrals can be evaluated for specified upper and lower values of *y* by using the series expansions in Equations (A3). The simplest way to obtain the corresponding conditional mean times to fixation and loss, *t*\* and *t*\*\*, for a new mutation with a haploid population size *N*_*H*_ (see next section) is to sum the products *t*\*(*x, q*) /*N*_*H*_ and *t*\*\*(*x, q*)/*N*_*H*_ over all values of *x* between *q* and 1, with *q* = 1/*N*_*H*_ (Ewens 2004, p.142).

## Theoretical results

### Approximate expected times to fixation and loss of a new mutation

A more general model of selection than that used by Mafessoni and Lachmann (2015) is employed here, using the notation of Charlesworth (2020a,b). A biallelic locus in a discrete generation, panmictic population is assumed, with frequencies *x* and 1 – *x* in a given generation of alleles A_2_ and A_1_, respectively; the population is initially fixed for A_1_. The new mutation, A_2_, is introduced as a single copy. To accommodate a general genetic system, the number of haploid genomes at a locus that are present in the population is denoted by *N*_*H*_, so that the initial frequency of A_2_ is *q* = 1/*N*_*H*_ (Charlesworth 2020a). Selection is sufficiently weak that second-order terms in the selection coefficient *s* for A_2_A_2_ homozygotes can be neglected (*s* is negative if A_2_A_2_ individuals are at a selective disadvantage); 1 + *s* is the fitness of A_2_A_2_ relative to the fitness of A_1_A_1_. The effective population size is *N*_*e*_; time is measured in units of the coalescent time, 2*N*_*e*_ generations. The scaled selection coefficient is defined as *γ* = 2*N*_*e*_*s*. Under these assumptions, the rate of change in frequency of A_2_ can be written as:

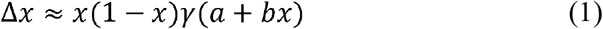

where the coefficients *a* and *b* depend on the genetic system and breeding system, and higher-order terms in *s* have been neglected (Charlesworth 2020a). In the case of an autosomal locus with random mating, *a* = *h* and *b* = 1 – 2*h*. Here, *h* is the dominance coefficient, such that the fitness of A_1_A_2_ heterozygotes relative to that of A_1_A_1_ homozygotes is equal to 1 + *hs*. This familiar case will be the focus of this study, but the conclusions can easily be extended to other cases, as described in Table 1 of Charlesworth (2020a). For example, for an autosomal locus and an inbreeding coefficient *F*, we have *a* = *F* + (1 – *F*)*h*, and *b* = (1 – *F*)(1 – 2*h*). The sign of *b* describes whether A_2_ is (partially) dominant or recessive, *b* being negative with dominance and positive with recessivity; *b* = 0 for a semi-dominant mutation.

**Table 1.** Parameters used in the model of associative overdominance.

|  |  |  |
| --- | --- | --- |
| 1019 | $u$ | rate of mutation per generation from $A_1$ to $A_2$ at the selected locus |
| 1020 | $v$ | rate of mutation per generation from $A_2$ to $A_1$ at the selected locus |
| 1021 | $\kappa$ | mutational bias parameter, such that $u = \kappa v$ |
| 1022 | $\theta_s$ | scaled mutation rate at the selected locus, such that $\theta_s = 4N_e u$ |
| 1023 | $P_{10}$ | Probability of loss of an $A_2$ mutation introduced into a population fixed for $A_1$. |
| 1024 | $P_{11}$ | Probability of fixation of an $A_2$ mutation introduced into a population fixed for $A_1$. |
| 1025 | $P_{20}$ | Probability of loss of an $A_1$ mutation introduced into a population fixed for $A_2$. |
| 1026 | $P_{21}$ | Probability of fixation of an $A_1$ mutation introduced into a population fixed for $A_2$. |
| 1027 | $\lambda_{10}$ | Rate of loss of new $A_2$ mutations from a population fixed for $A_1$ (in units of $2N_e$ |
| 1028 | | generations); $\lambda_{10} = 0.5 \theta_s N_H P_{10}$ |
| 1029 | $\lambda_{11}$ | Rate of fixation of new $A_2$ mutations in a population fixed for $A_1$ (in units of $2N_e$ |
| 1030 | | generations); $\lambda_{11} = 0.5 \theta_s N_H P_{11}$ |
| 1031 | $\lambda_{20}$ | Rate of loss of new $A_1$ mutations from a population fixed for $A_2$ (in units of $2N_e$ |
| 1032 | | generations); $\lambda_{20} = 0.5 \kappa^{-1} \theta_s N_H P_{20}$ |
| 1033 | $\lambda_{21}$ | Rate of fixation of new $A_1$ mutations in a population fixed for $A_2$ (in units of $2N_e$ |
| 1034 | | generations); $\lambda_{21} = 0.5 \kappa^{-1} \theta_s N_H P_{21}$ |
| 1035 | $\Delta\pi_{il}$ | Value of $\Delta\pi$ at the neutral site, when an allele of type $A_i$ has just become lost from the |
| 1036 |  | linked selected site. |
| 1037 | $\Delta\pi_{if}$ | Value of $\Delta\pi$ at the neutral site, when an allele of type $A_i$ has just become fixed at the |
| 1038 |  | linked selected site. |
| 1039 | $\Delta\pi_{i0s}$ | Sum of $\Delta\pi$ values at the neutral site over the course of the loss of a type $i$ mutation from the |
| 1040 |  | linked selected site. |
| 1041 | $\Delta\pi_{i1s}$ | Sum of $\Delta\pi$ values at the neutral site over the course of the fixation of a type $i$ mutation at the |
| 1042 |  | linked selected site. |
| 1043 |  |  |

For the present purpose, the main quantities of interest are the sojourn time densities *t*\*(*x, q*) and *t*\*\*(*x, q*) for mutations destined for fixation and loss, respectively, as defined in the Material and Methods, *Numerical integrations*. These are expressed in units of the coalescent time 2*N*_*e*_; the corresponding absolute times can be obtained by multiplying by 2*N*_*e*_. When A_2_ is initially present as a single copy, only the situation when *q* ≤ *x* ≤ 1 need be considered; the relevant expressions for *t*\*(*x, q*) and *t*\*\*(*x, q*) are given in the Appendix. Approximations that use only first- and second-order terms in *γ* are given by Equations (A5) and (A7), for *t*\*(*x*, 1/*N*_*H*_) and *t*\*\*(*x*, 1/*N*_*H*_), respectively.

Equations (A5) and (A7) can be integrated over the interval 1/*N*_*H*_ ≤ *x* ≤ 1, yielding approximate expected conditional times to fixation and loss for a new mutation. We have:

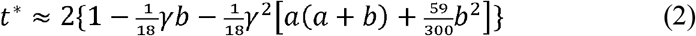

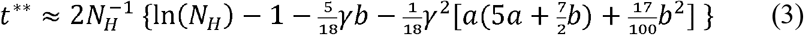

For greater accuracy, Euler’s constant (0.5772..) should be added to the terms inside the brackets on the right-hand side of Equation (3), to correct for the difference between the summation and integration of 1/*x* (Ewens 2004, p.23). Kaushik and Jain (2021, Equation 9) have independently derived an approximation similar to Equation (2) for the case of an autosomal locus and random mating. Their *H* corresponds to *h* – ½; if terms in *H*^2^ are neglected, their equation is the same as Equation (2), except for the fact that their expression for *t*\* differs by a factor of ½, because of their assumption of a continuous time birth-death process rather than a Wright-Fisher model.

The first-order terms in *γ* in these equations show that when *b* > 0 (*h* < 0.5 in the case of autosomal inheritance), the mean times to fixation and loss of a favorable mutation (*γ* > 0) are reduced below their neutral value, if *γ* is sufficiently close to 0. These times are increased when *b <* 0 (*h* > 0.5 with autosomal inheritance). The converse relations hold for the case of a deleterious mutation (*γ* < 0).

These results correspond to those of Mafessoni and Lachmann (2015), based on numerical evaluations of the relevant general equations. Furthermore, when *b* = 0 (corresponding to semi-dominance with diploid inheritance), *t*\* and *t*\*\* are at a maximum with respect to *γ* when *γ* = 0, so that selection on a semi-dominant autosomal mutation (or a semi-dominant mutation with respect to female fitness with X-linkage) is always associated with a shorter conditional time to fixation or loss than under neutrality. The results are of interest for the main topic of this paper, because the flux of mutations between deleterious and favorable alleles at a site is the basis for the analysis of the long-term effects of weak selection on variability (see the section below on *Relevance to associative overdominance and background selection*). An analysis of the effects of the quadratic terms in *γ* on the implications of Equations (2) and (3) is given in the Supplementary File S1, section 1.

A similar treatment can be given for the means of the sum of the diversity 2*x*(1 – *x*) in each generation over the paths to fixation or loss of a new mutation (*H\** and *H\*\**, respectively), which are obtained by integrating the product of 4*N*_*e*_ *x*(1 – *x*) with *t*\*(*x, q*) or *t*\*\*(*x, q*), respectively, over *x* between 0 and 1. The relevant integrations give:

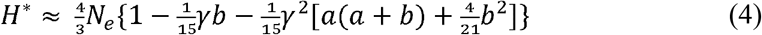

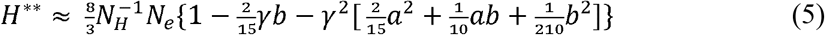

These expressions show that *H\** and *H\*\** have similar properties to *t\** and *t\*\** with respect to their dependence on *b* when selection is sufficiently weak that first-order terms in *γ* predominate. Weak selection with *γb* < 0 can thus result in an increase in these measures of diversity relative to neutral expectation, contrary to what is commonly assumed in discussions of molecular variability. This can happen either when *γ* < 0 and *b* > 0 (a deleterious, partially recessive or recessive mutation) or *γ* < 0 and *b* > 0 (a favorable, partially dominant or dominant mutation).

The sojourn time density functions in Equations (A8) and (A9) can be used to find the corresponding net mean sojourn time between loss or fixation for a new mutation, to an accuracy of order *γ* ^2^:

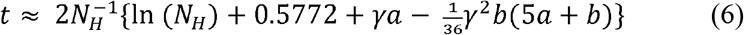

Similarly, the expected sum of the diversity values over the sojourn of a mutation in the population before its loss or fixation is approximated by:

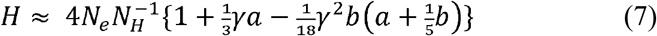

With respect to terms of the first order in *γ*, there is no dependence on *γb* of the net mean sojourn time and net diversity, but they are both increasing functions of *γa*. This is consistent with the classical result for the case of a semi-dominant autosomal mutation with random mating, where *H* is an increasing function of *γ* (Fisher 1930, Figure 3; Kimura 1983, p.44). At first sight, it seems paradoxical that semi-dominant mutations subject to positive selection should yield a higher net diversity than neutral mutations during their sojourn in the population. But this is simply a reflection of the fact that they have a higher chance of establishing themselves in the population, and hence of contributing to diversity.

### Numerical results for conditional times to loss and fixation and net diversity

The accuracy of these approximations was checked by comparison with the results of numerical integrations of the relevant diffusion equation formulae, as described in the Material and Methods and the Appendix, and by computer simulations of an autosomal locus in a randomly mating population of size *N*, with binomial sampling of post-selection gametes. In this case, *N*_*e*_ = *N* and *N*_*H*_ = 2*N*. Figure 1 shows the mean fixation time of a new mutation (conditioned on its fixation) for a range of values of the magnitude of the scaled selection coefficient, |*γ*| = 2*N*_*e*_ |*s*|. Both deleterious mutations (*s* < 0) with dominance coefficient *h* = 0.1 and favorable mutations with dominance coefficient *h* = 0.9 (*s* > 0) were modeled. The values for a neutral mutation, obtained by integration of the sojourn time density function in the absence of selection with respect to *x* between 1/ *N*_*H*_ and 1 – 1/ *N*_*H*_, are indicated by the horizontal lines. The results from the numerical integrations of the sojourn time densities (black curves) are in close agreement with the simulation results (red points) over the whole range of *γ* values used here. The approximate values from Equation (2) (blue curves) are in good agreement with the more exact values for |*γ*| up to 2 or so, but then tend to overestimate the fixation times. Nonetheless, the qualitative pattern of an increase in fixation time above the neutral value as |*γ*| increases to a value near 2, followed by a decrease, is captured by the approximation. The more exact results show that the neutral value of *t*\* is returned to more quickly, at |*γ*| ≈ 3, than is indicated by the approximation. As expected from the results of Maruyama (1972) and Maruyama and Kimura (1974), the curves for favorable and deleterious mutations, with their complementary values of *h*, are identical.

**Figure 1.**
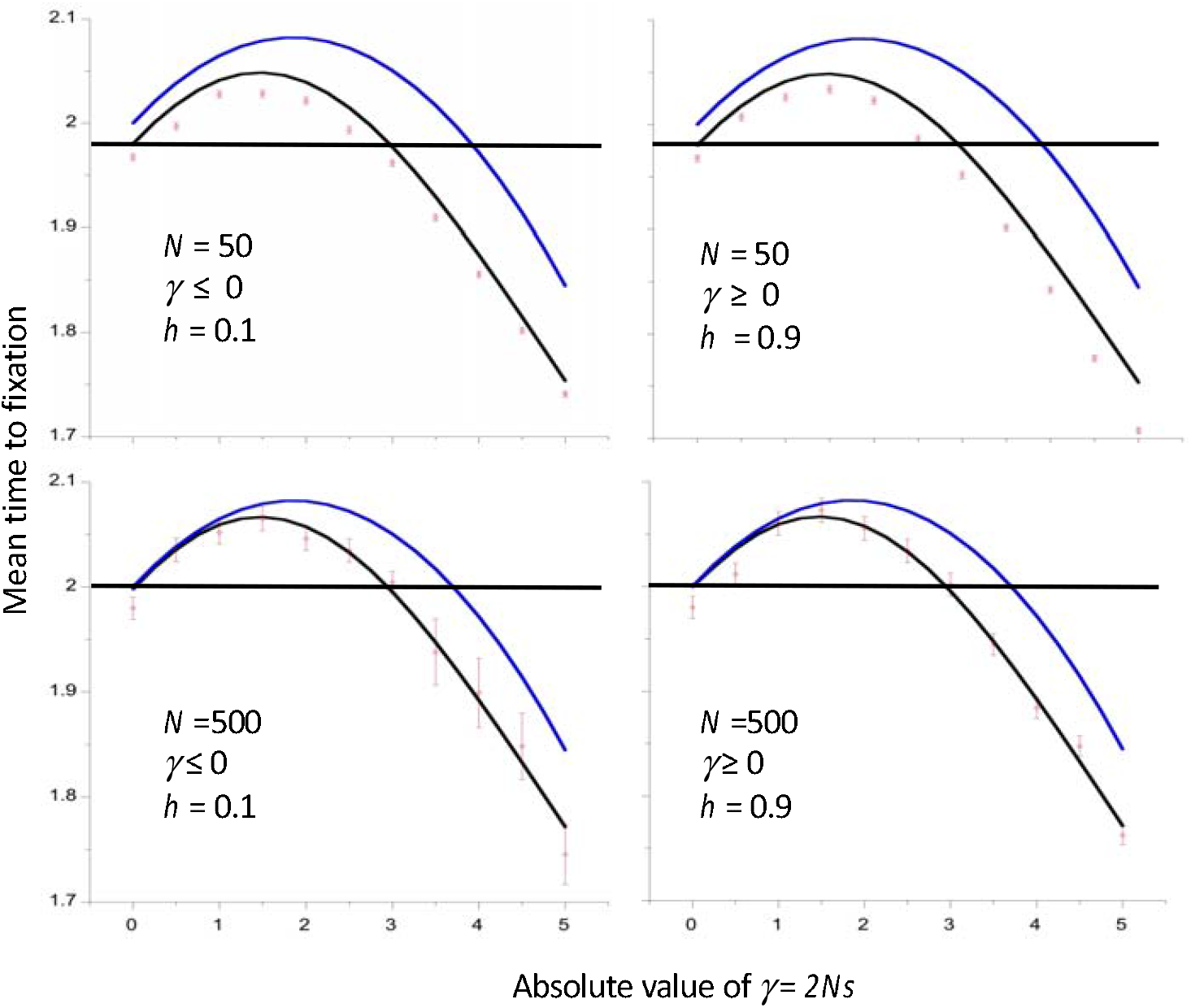
Mean time to fixation of a new mutation as a function of the absolute value of the scaled selection coefficient *γ*, assuming autosomal inheritance and a Wright-Fisher model of genetic drift with population size *N*. Times are in units of coalescent time (2*N* generations). The cases with negative selection have a dominance coefficient *h* = 0.1 and those with positive selection have *h* = 0.9. The results for two different values of *N* are shown. The red points with error bars are the mean values from simulations and their standard errors. The blue curves are the values from the approximation of Equation (2), and the black curves are the values from numerical integrations of the diffusion equation results (Equations A1). The horizontal lines are the neutral values from the numerical integrations.

Figure 2 shows similar results for the mean times to loss of new mutations, conditioned on their loss. These are necessarily much smaller than the mean times to fixation, and are also inverse functions of the population size when measured on the coalescent timescale of 2*N*_*e*_ generations, as implied by Equation (3). In terms of generations, the loss times are logarithmically increasing functions of *N*, since *N*_*e*_ = *N* in the cases shown here. There are noticeable differences between the results for deleterious and favorable mutations, with the partially deleterious mutations having longer mean times to loss than the complementary partially dominant favorable mutations (with dominance coefficient equal to 1 – *h*, where *h* is the value for the corresponding partially recessive mutations), due to the multiplicands of –*γ*^2^ being smaller for the deleterious than the complementary favorable mutations. Similarly, the deleterious mutations return to the neutral value at larger values of |*γ*| than the complementary favorable mutations. The multiplicand of –*γ b* is larger for losses than fixations, so that the condition on |*γ*| for *t*\*\* to exceed the neutral value is more liberal for losses than for fixations. However, the second-order terms in both cases are always positive, and overcome the first-order terms when |*γ*| is large enough.

**Figure 2.**
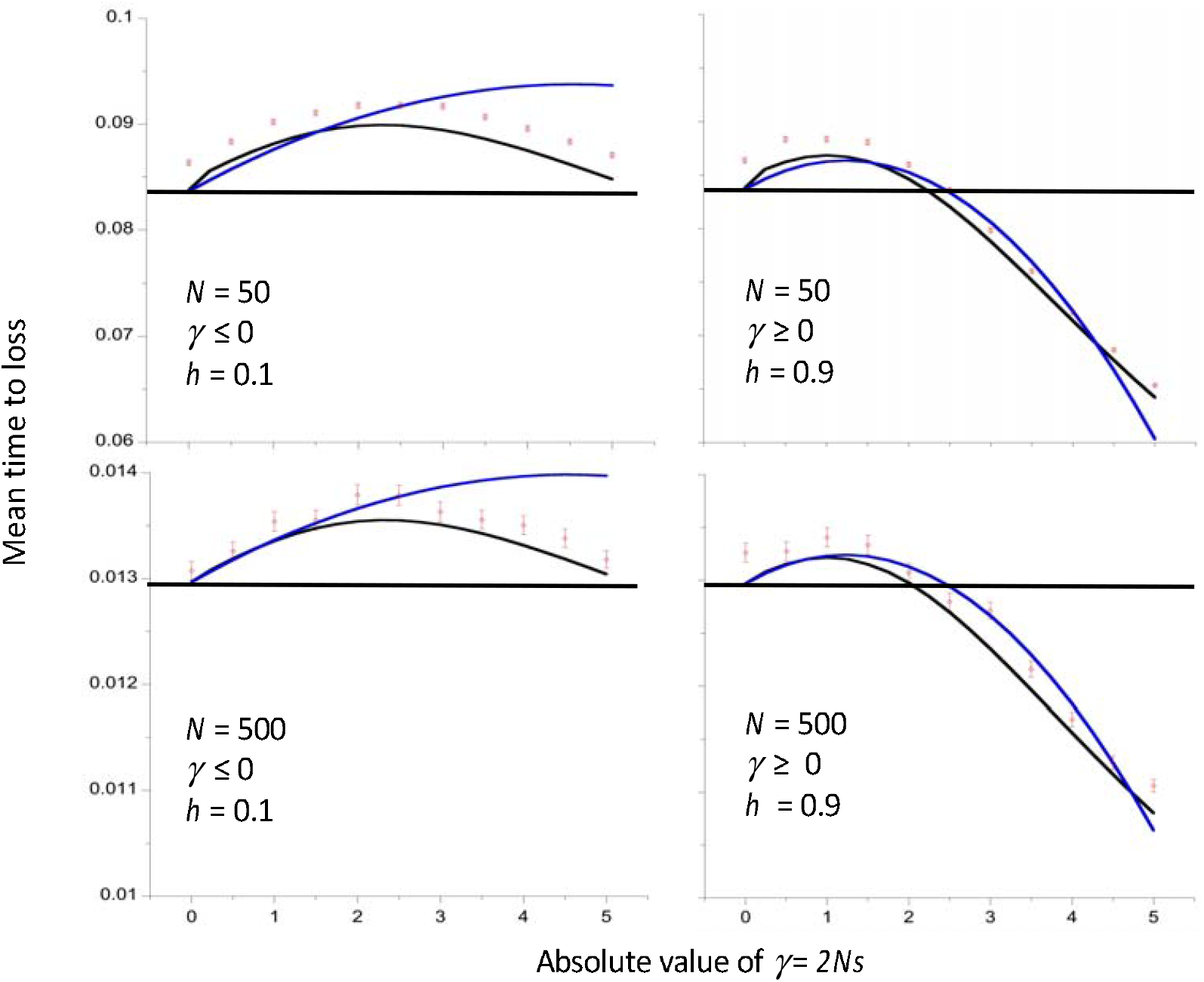
Mean time to loss of a new mutation as a function of the absolute value of the scaled selection coefficient, assuming autosomal inheritance and a Wright-Fisher model of genetic drift with population size *N*. Times are in units of coalescent time (2*N* generations). The cases with negative selection have a dominance coefficient *h* = 0.1 and those with positive selection have *h* = 0.9. The results for two different values of the population size, *N*, are shown. The red points with error bars show the mean values of simulations and their standard errors. The blue curves are the values from the approximation of Equation (3), and the black curves are the values from numerical integrations of the diffusion equation results (Equations A1 and A6). The horizontal lines are the neutral values from the numerical integrations.

Similar patterns of behavior are found for the mean sums of the diversities conditioned on fixations or losses, *H*\* and *H*\*\*; examples similar to those in Figures 1 and 2 are shown in Figures S1 and S2, using the integration and approximate results. As might be expected, *H*\* and *H*\*\*are closely correlated with *t*\* and *t*\*\*, respectively (Figure S3).

Figure 3 shows the approximate and integration results for the conditional mean times to fixation and loss for the whole range of dominance coefficients, with the magnitude of the scaled selection coefficient |*γ*| equal to 1 or 2. The approximations (solid curves) agree very well with the integration results (dashed curves) for both strengths of selection, and are nearly linear in *h*. As expected from Equations 2 and 3, the relationships of the fixation and loss times to *h* for the deleterious and favorable mutations are opposite in direction, with deleterious mutations having fixation and loss times that decrease with *h*, whereas those for favorable mutations increase with *h*. Weaker selection is associated with a larger range of *h* values for which the conditional fixation time is greater than the neutral value.

**Figure 3.**
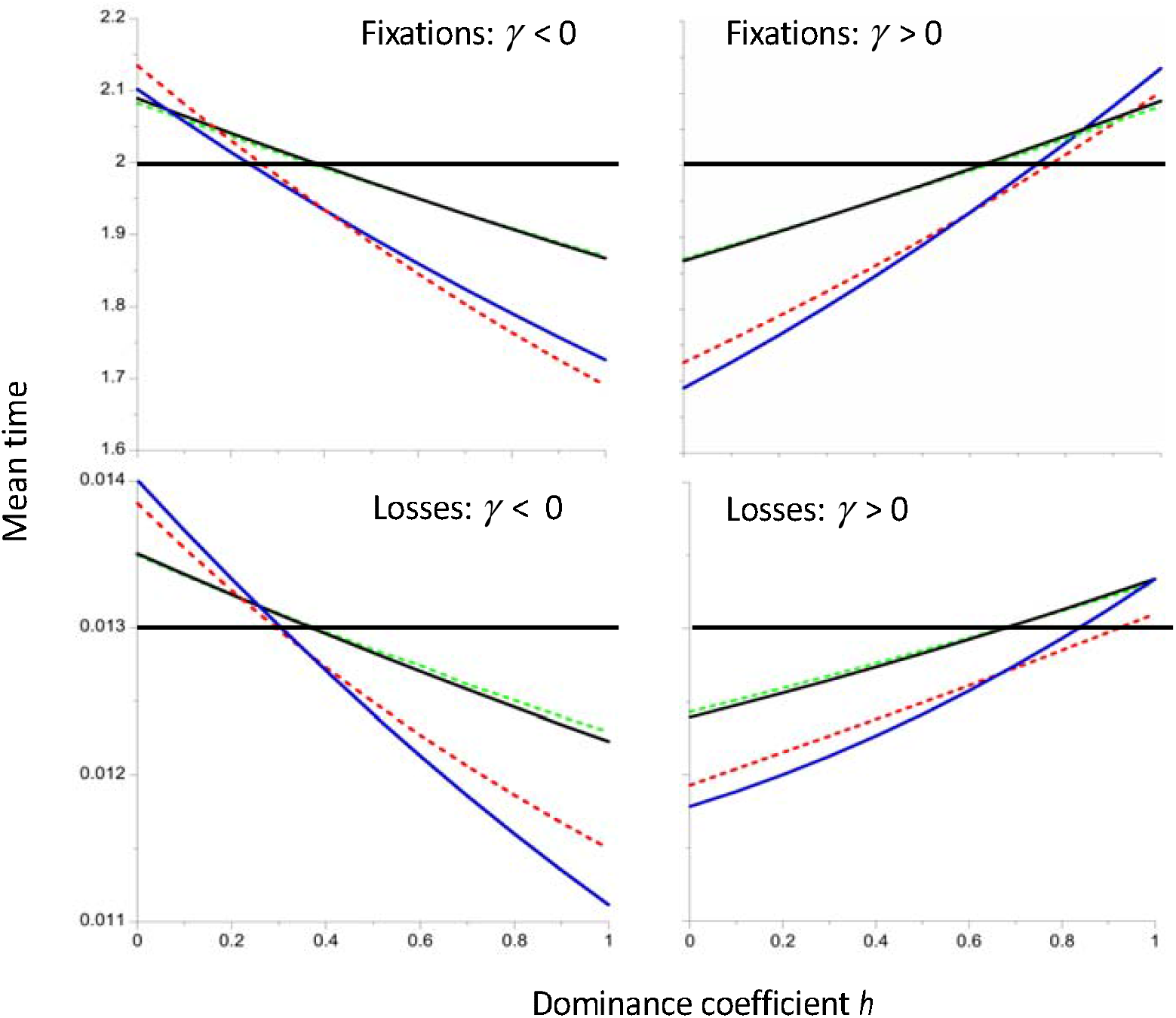
Mean time to fixation and loss of a new mutation (in units of coalescent time) as a function of the dominance coefficient *h*, for the models used in Figures 1 and 2. The black and blue lines use the approximations of Equations (2) and (3) to generate results for |*γ*| = 1 and 2, respectively. The dashed green and red lines are the corresponding values from numerical integrations of the diffusion equation expressions. The horizontal lines are the neutral values from the numerical integrations. The population size *N* is equal to 500.

A critical value of the dominance coefficient for a given *γ, h*_*cf*_ (*γ*), can be defined, which is the value of *h* at which the mean fixation time is equal to the neutral value. A similar quantity, *h*_*cl*_(*γ*), can be defined for the mean time to loss. As is discussed in more detail in the section *Relevance to associative overdominance and background selection*, these quantities provide a useful link between the present approach and that of Zhao and Charlesworth (2016), who examined the conditions for AOD versus BGS for the case of a large population that is initially in stochastic equilibrium under mutation, drift and selection, and then suffers a permanent reduction in population size. As a measure of the magnitude of genetic drift, they used the asymptotic rate at which variability at a neutral site linked to a site under selection decreases after the reduction in *N*, and determined whether selection increased or reduced the rate of loss of variability.

Approximations for the critical dominance coefficients that correspond to the boundary between an increased and a decreased mean fixation or loss time can be obtained by setting the sum of the terms involving *γ* and *γ*^2^ to zero in Equations (2) and (3), and solving the resulting quadratic equations in *h*. For *h*_*cf*_ (*γ*), the further approximation of replacing 59/300 with 1/5 is used.

After some algebra, the following expressions are obtained:

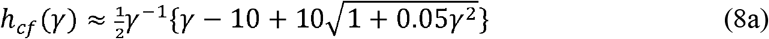

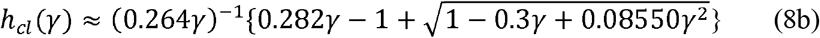

As *γ* approaches zero, consideration of the leading terms in *γ* in the radicals in these equations shows that the critical dominance coefficient ≈ 0.5 + 0.125*γ* in both cases, so that slightly recessive mutations (*γ* < 0) or dominant mutations (*γ* > 0) result in an increase in mean fixation and loss times when selection is very weak. For small *γ*, which is the main region of interest as far as AOD is concerned, the critical dominance coefficients are nearly linear in *γ* (Figure 4).

**Figure 4.**
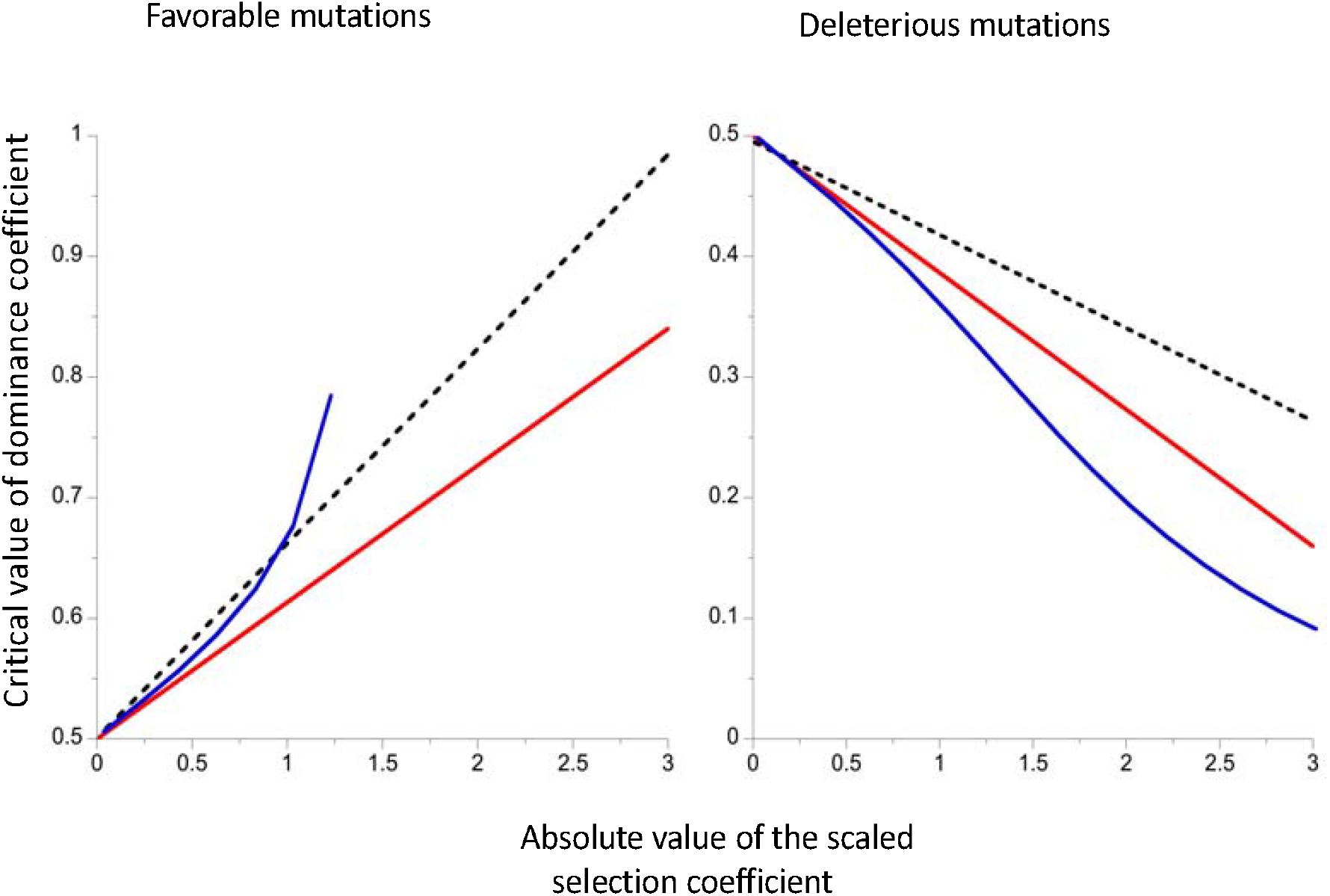
The critical values of the dominance coefficient as functions of the scaled selection coefficient for mean fixation time (full red line) or mean loss time (dashed black line). The results were obtained using the approximate Equations (8). The blue curves are the solutions to Equation (18) of Zhao and Charlesworth (2016). The results for favorable mutations are shown in the left-hand panel) and those for deleterious mutations in the right-hand panel

The blue curves in Figure 4 show the critical dominance coefficients for the operation of AOD, given by the solution of the approximate cubic Equation (18) of Zhao and Charlesworth (2016). For favorable mutations, this expression breaks down for *γ* > 1.26, but performs well for deleterious mutations over the entire range displayed. If the cubic equation is approximated by ignoring terms in *γ*^2^, the critical dominance coefficient for the operation of AOD is 0.5 + 0.125*γ*, the same as for *h*_*cf*_ and *h*_*cl*_. This suggests that the two approaches are related to each other, although the three different functions describing *h*_*c*_ do not agree precisely.

### Effects of weak selection on variability at linked neutral sites

Intuitively, one would expect the longer mean fixation times associated with weak selection and *γb* < 0 to produce a higher level of mean pairwise diversity (*π*) at a linked neutral site than when a neutral mutation has become fixed. This was shown to be the case by Mafessoni and Lachmann (2015) and Johri *et al*. (2021), using computer simulations (similar, but smaller, effects would be expected from losses of new mutations; these were not, however, examined in these papers). Note, however, that the fixation of a neutral mutation produces an approximately 42% reduction in *π* at completely linked sites (Tajima 1990), so that all these cases are still associated with reduced rather than increased diversity. This is because *π* is measured at a single point in time (when a new mutation has been fixed), rather than using an estimate of the mean of *π* over a long time period at a focal neutral site that is subject to a succession of evolutionary events at a linked site – if this site is neutral, the mean *π* at the focal site cannot be affected by these events. An approximate method for calculating *π* at the end of a fixation event was used by Johri *et al*. (2021), based on the selective sweep equations of Charlesworth (2020b); this procedure is, however, unsatisfactory, since the sweep equations are inaccurate at small |*γ*| values.

A first step towards obtaining an estimate of the long term mean value of *π* is to determine the pattern of variability at a focal neutral site over the course of fixation (or loss) of a linked mutation (A_2_) introduced into a population initially fixed for the alternative allele (A_1_). This was done here by applying the simulation procedure of Tajima (1990), as described in the Material and Methods section and in Johri *et al*. (2021). The left-hand panel in Figure 5 shows the results for fixations of a favorable mutation with dominance coefficient *h* = 0.9 and a range of values of the scaled selection coefficient, *γ*, in a population size of 50, and with complete linkage between the neutral and selected sites. All diversity statistics are measured relative to *θ* = 4*N*_*e*_*µ*, the equilibrium value in the absence of selection. It will be seen that the mean relative *π* over all genotypes taken over a fixation event (*π*_*w*_) exceeds one for sufficiently small *γ* values, even for the neutral case of *γ* = 0, whereas the relative *π* at the time of fixation or loss is always < 1. The reason for the excess in *π*_*w*_ over neutral expectation is that the slowness of the fixation process (whose expected value is of the order of 4*N*_*e*_ generations with weak selection) allows A_1_ and A_2_ haplotypes to diverge in sequence over the course of the fixation of A_2_, because their mean coalescent times in the absence of recombination are much greater than the standard neutral value of 4*N*_*e*_ generations (Kimura and Ohta 1969). Comparable results are seen for losses of new mutations, but with much smaller effects due to the shorter expected duration of loss events, which is close to 2 ln(*N*_*H*_) generations with weak selection (Figure S4).

**Figure 5.**
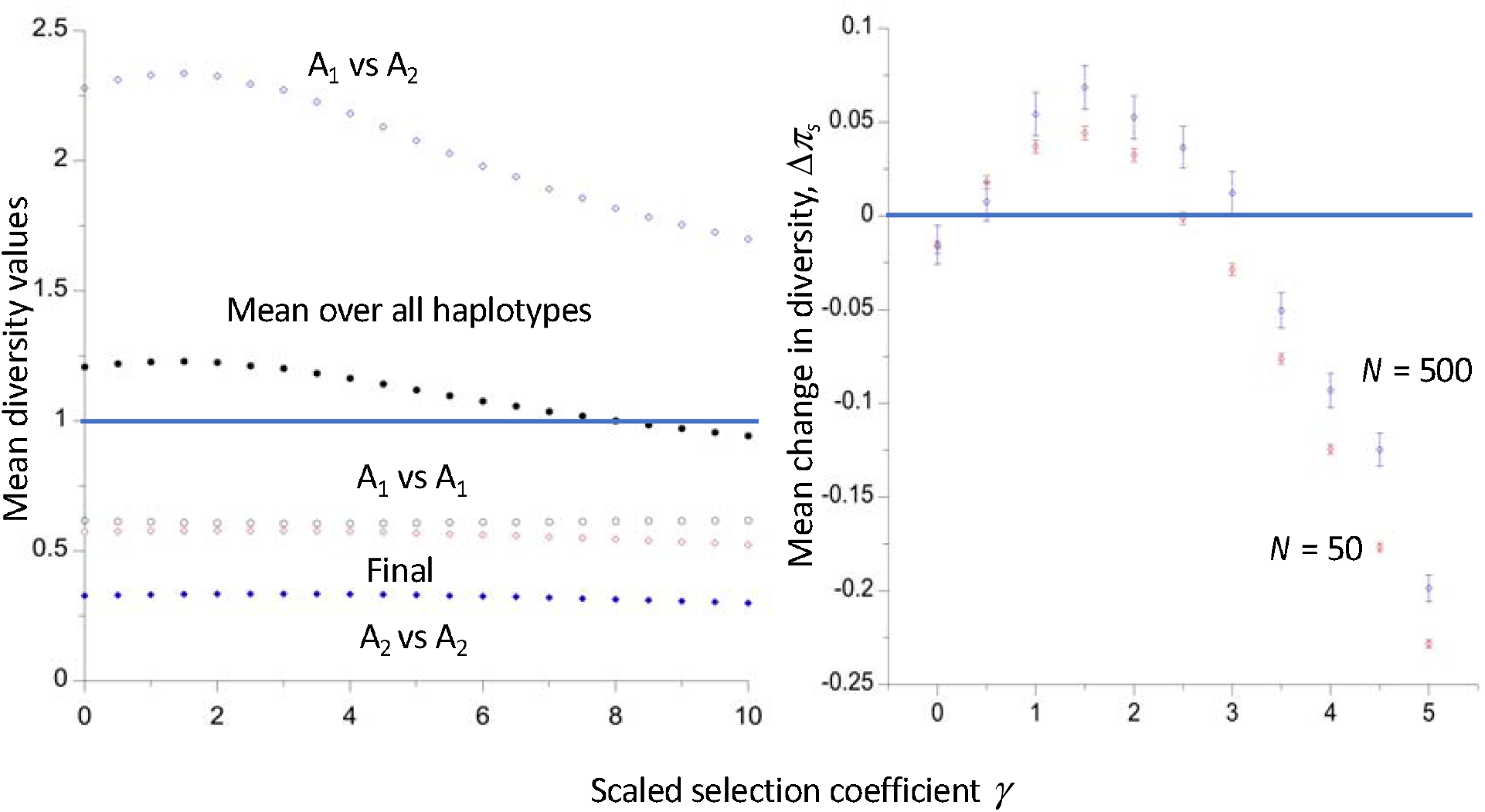
The left-hand panel shows the mean pairwise diversities (relative to the purely neutral value *θ*) at a neutral site during the course of fixation of a completely linked beneficial mutation with dominance coefficient *h* = 0.9, for haplotypes carrying the ancestral allele (A_1_), the new mutation (A_2_), and the divergence between A_1_ and A_2_ haplotypes (A_1_ versus A_2_). The final diversity at the time of fixation of A_2,_ and the mean diversity over all three haplotypes over the course of the fixation process, are also shown. Autosomal inheritance and a Wright-Fisher model of genetic drift with population size *N* = 50 were simulated by the method of Tajima (1990). The right-hand panel displays the mean values and standard errors of the net change in relative diversity over repeated fixation events for two different *N* values, estimated from the simulations by the method described in the text.

These results yield the seemingly paradoxical conclusion that fixation and loss events can be associated with a net increase in mean *π* at linked neutral sites during the course of fixation or loss, even when they involve neutral mutations. The paradox with respect to neutral mutations comes from the fact that one type of conditional event has been replaced by another: we are conditioning on observing a fixation or loss event that is in progress, and the increasing divergence between haplotypes carrying the ancestral and mutant alleles influences the net effect of the fixation or loss event on variability at the neutral site, as discussed in more detail below.

The following approximate approach avoids such conditioning. As was done by Charlesworth (2020b) for the case of sweeps of strongly selected mutations, ergodicity is assumed: the probability of a given value of *π* resulting from the hitchhiking effects of the selected site is proportional to the amount of time that the process spends at that value of *π*, analogous to the use of the sojourn time density for determining the frequency spectrum at a single locus (Ewens 2004 pp.24-25).

A core assumption is that the rates of fixation and loss events are sufficiently low that the intervals between successive events allow a complete recovery of diversity. Once again, diversities are measured relative to the value in the absence of selection, and the deviation of the relative diversity value from one is denoted by Δ*π*. If the mean value of Δ*π* immediately after a given fixation or loss event is denoted by Δ*π*_0_, and the value of Δ*π* at time *t* after such an event is Δ*π*_*t*_, we have Δ*π*_*t*_ ≈ Δ*π*_0_ exp(– *t*), where *t* is in units of 2*N*_*e*_ generations (Malécot 1969, p.40; Wiehe and Stephan 1993, Equation 6a). The sum of the values of Δ*π*_*t*_ over subsequent generations, relative to 2*N*_*e*_, can thus be approximated by:

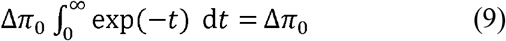

We also need to consider the contribution from the mean diversity over the course of each fixation (loss) event, which is given by the expectation of the product of the duration of an event, *T*_*e*_, and the associated mean value per generation of Δ*π* over the course of the event, Δ*π*_*w*_, *i*.*e*., by E{*T*_*e*_Δ*π*_*w*_}. If *T*_*e*_ is measured in units of coalescent time, and we use the sum of the Δ*π* values relative to 2*N*_*e*_ over a long period of time (or over many independent pieces of genome that are all subject to the same evolutionary process), the mean sum of the deviation from 1 of the relative diversity between successive fixation (loss) events is given by:

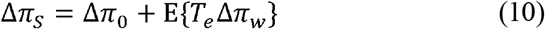

To calculate the change in mean relative diversity due to a succession of evolutionary events, we would of course need to know the total length of time involved, requiring a calculation of the expected number of occurrences of fixation (loss) events and the expected times between them (see the next section). Use of Equation (10) avoids having to make a detailed model of these events, and provides an index of the expected magnitude and sign of the effects on neutral diversity of fixations (losses) at a linked site. Δ*π*_*S*_ can thus be (loosely) referred to as the mean change in diversity associated with a series of fixation or loss events.

It should be noted that the assumption of an indefinitely large amount of time for recovery of diversity between fixation or loss events implies that the contribution of Δ*π*_0_ relative to E{*T*_*e*_Δ*π*_*w*_} is overestimated compared with a realistic evolutionary model, so that any increases in diversity with weak selection when *γb* < 0 will be underestimated by this approach when Δ*π*_0_ < 0 and E{*T*_*e*_Δ*π*_*w*_} > 0.

Numerical values of the quantities involved can be obtained from the simulations of a neutral sites linked to a locus subject to selection, using the algorithm of Tajima (1990), as described in the Material and Methods. The relevant statistics are shown in the left-hand panel of Figure 5 for the case of the fixation of a partially dominant, favorable mutation (*h* = 0.9) in a population with *N* = 50. Using Equation (10), Δ*π*_*S*_ for recurrent fixations is shown in the right-hand panel of Figure 5, for *N* = 50 and *N* = 500. Here, the mean of *T*_*e*_Δπ_*w*_ over many replicate simulations is used as an estimate of the expectation of *T*_*e*_Δπ_*w*_.

It can be seen that fixations of highly dominant, favorable mutations cause an increase in diversity when *γ* is sufficiently small, but diversity is reduced when *γ* is somewhat greater than 2; the value of *γ* at the point of return to a reduction in *π* is smaller with the larger value of *N*. As expected from the properties of the mean fixation time described above, a similar pattern (within the limits of statistical error) is observed for fixations of deleterious mutations with *h* = 0.1 and the same absolute value of *γ* (Figure S4). Figures 6 and S5 show comparable results for losses of both favorable and deleterious mutations; in this case, the patterns are noticeably different for the two types of mutations, and are much more strongly affected by the population size, as expected from the corresponding effects on mean loss times described in the previous section. Tables S1-S8 show the detailed statistics for the simulations on which these figures are based, and Table S9 shows summary diversity statistics for fixation and loss events for a range of dominance coefficients, and two different strengths of positive and negative selection.

**Figure 6.**
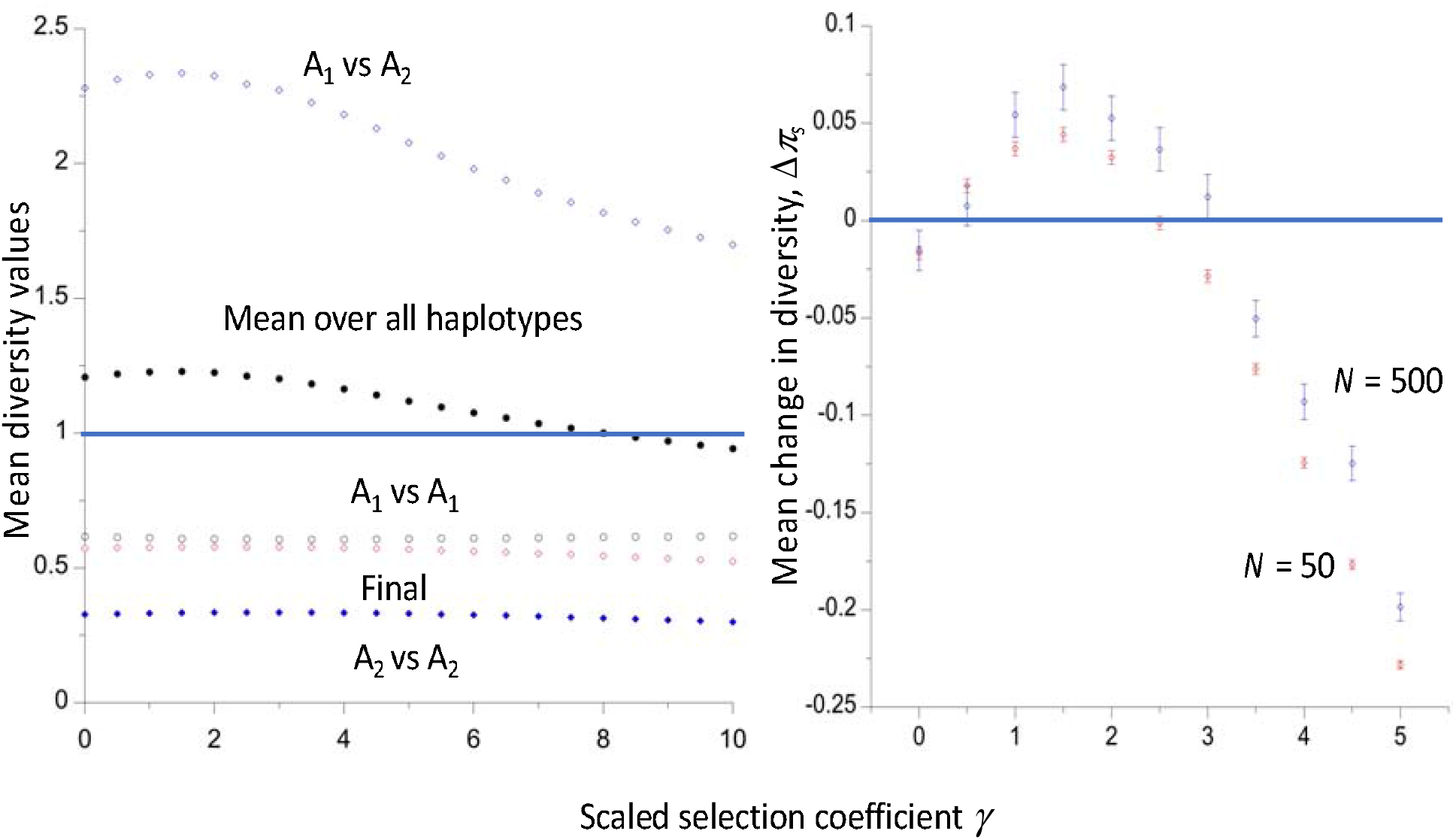
The mean values and standard errors of the net changes in relative diversity over repeated loss events with two different *N* values, for deleterious mutations with *h* = 0.1 and favorable mutations with *h* = 0.9, using the same model and methods as in Figure 7.

The results allow questions to be asked about the nature of the major determinant of the effect of selection on neutral diversity at a linked site. Intuitively, it would seem likely that the mean conditional times to fixation, *t*\*, and loss, *t*\*\*, must play a major role. However, the answer depends on what aspect of diversity is being explored. A detailed discussion of results relating to this question is given in section 2 of Supplementary File 1, section 1. These results show that, for a given value of *γ*, both sojourn times are nearly linearly related to the diversity statistics, but the sign and magnitude of *γ* influence the values of the diversity statistics for the same value of *t*\* or *t*\*\*. There is a close correspondence between the values of *t*\* and *t*\*\* that are generated by the simulations with *γ* = 0 and the thresholds at which Δ*π*_*S*_ ≥ 0, within the limits of sampling error (Figures S6 and S9).

The results can be interpreted as follows. The mean neutral diversity of haplotypes carrying the selectively favorable A_2_ variant increases with the duration of a fixation event, because these haplotypes spend longer in the population before reaching fixation and hence have a longer coalescence time. The divergence between A_1_ and A_2_ haplotypes also increases with *t*\*, since a longer time is available for them to diverge. The changes in the diversity of A_1_ haplotypes are more minor. The net result is that the net mean diversity over the course of the event (Δ*π*_*w*_) increases with *t*\*, as can be seen in Figure S7, so that its product with *t*\*, which appears in the equation for Δ*π*_*S*_ (Equation 10) also increases with *t*\*. This effect is reinforced by the decline in the reduction of diversity at the time of fixation of A_2_, – Δ*π*_0_, as *t*\* increases (Figure S8), reflecting a greater coalescence time for A_2_ when its sojourn time is longer.

These considerations apply to both positive and negative selection. However, with positive selection, A_2_ spends more of its time at high frequencies than at low frequencies before becoming fixed, whereas the reverse is true with negative selection, as can be seen in the plots of the sojourn time densities relative to the neutral values that are shown in Figure 8 (note that *t*\*(*x*) is independent of *x* in the absence of selection). This means that, for the same |*γ*| value, haplotypes carrying A_1_ have a longer coalescence time with negative than with positive selection; because these haplotypes start with the equilibrium neutral diversity value *θ*, they contribute disproportionately to Δ*π*_*w*_. This effect is especially clear for the larger values of |*γ*|; for example, with |*γ*| = 5, the mean diversity of A_1_ haplotypes is 0.657±0.001 for *γ* = –5, and 0.608±0.001 for *γ* = 5.

Similar patterns are seen with losses of mutations. This is perhaps less surprising, since *t*\*\* for a given |*γ*| value is greater for deleterious mutations than for favorable mutations when *h* for the former is the same as 1 – *h* for the latter, i.e., the dominance coefficients are complementary (Figure 3).

### Relevance to associative overdominance and background selection

As mentioned in the Introduction, the relevance of these results to associative overdominance (AOD) arises from the fact that, if selection is sufficiently weak in relation to genetic drift, a biallelic locus subject to selection and reversible mutations will experience a constant flux of mutations from the states of fixation for A_1_ to fixation for A_2_ and vice-versa, as in the Li-Bulmer model of codon usage bias (Li 1987; Bulmer 1991; McVean and Charlesworth 1999). Here, A_1_ is now used to denote the selectively favorable allele at a given site, and A_2_ its deleterious alternative, rather than ancestral and mutant alleles, respectively.

For the case of an autosomal locus, the relative fitnesses of the three genotypes A_1_A_1_, A_1_A_2_ and A_2_A_2_ are 1, 1 – *hs* and 1 – *s*, respectively (*s* ≥ 0); with weak selection, this is equivalent to representing these fitnesses as 1 + *s*, 1 + (1 – *h*)*s* and 1. Thus, if the deleterious effects of A_2_ are recessive or partially recessive (0 ≤ *h* < 0.5) and *γ* = 2*N*_*e*_*s* is sufficiently small, the entry of an A_2_ mutation into a population temporarily fixed for A_1_ will be associated with a longer net sojourn time in the population than that for a neutral mutation. The same applies to the entry of a favorable A_1_ mutation into a population temporarily fixed for A_2_.

The results in the previous section strongly suggest that, under these conditions, diversity at closely linked neutral sites will be enhanced when a stationary state with respect to mutation, selection and drift is reached. The effect of the constant flux of mutations at a selected site on the expected level of neutral diversity (*π*) at the linked neutral site can be determined with the use of the ergodic assumption. This allows the application of the results of the simulations described above that used this approach, employing the infinite sites assumption that mutations are sufficiently infrequent that new mutations occur only at sites that are fixed for either A_1_ or A_2_, which was applied to the theory of codon usage by Bulmer (1991) and McVean and Charlesworth (1999).

Table 1 shows the variables that are used in this procedure. As in the previous section, diversity is measured relative to the purely neutral equilibrium value *θ =* 4*N*_*e*_*µ*, and its deviation from one at given point in time is denoted by Δ*π*.

Consider a site that has just become fixed for A_1_. Under the above assumptions, the distribution of times until an A_2_ mutation arises and becomes fixed is exponential, with parameter *λ*_11_. If a time *t* is drawn from this distribution, the expected number of A_2_ mutations that arise and become lost during an interval of length *t* is *λ*_10_ *t*, where *λ*_10_ >> *λ*_11_ from the formulae in Table 1. Integrating over the distribution of *t*, the expected number of losses of A_2_ mutations before A_2_ becomes fixed is found to be approximately equal to *λ*_10_ /*λ*_11_ = *P*_10_/*P*_11_. If the times between each successive event are >> 1, diversity completely recovers between loss events, and between the initial fixation and first loss event (it is shown below that this assumption can be relaxed). Applying Equation (10) from the previous section, and using the notation in Table 1, the expected sum (relative to 2*N*_*e*_) of Δ*π* values at the neutral site over the interval between the successive fixation events A_2_ to A_1_ and A_1_ to A_2_ is given by:

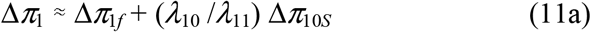

A similar argument applies to the interval succeeding the fixation of the A_2_ mutation and another fixation of an A_1_ mutation, substituting 2 for 1 in the first subscripts for the *P*s and *λ*s in Table 1 (*λ*_20_ >> *λ*_21_ is assumed), giving:

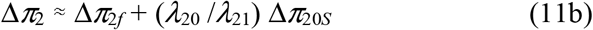

The expectation of the sum of Δ*π* values over an entire cycle between successive fixations of A_1_ mutations is simply Δ*π*_*T*_ = Δ*π*_1_ + Δ*π*_2_. We can thus estimate the expected value of Δ*π*_*T*_ per generation, Δ*π*_*e*_, by dividing Δ*π*_*T*_ by the expected time between successive fixations of A_1_ (in units of 2*N*_*e*_ generations), *T*_*s*_ = (*λ*_11_^−1^ + *λ*_21_^−1^):

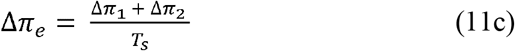

It is useful to note that the expression for *T*_*s*_ is consistent with the existence of a stationary state for the proportion of sites fixed for A_1_ versus A_2_, using the infinite sites assumption. If these proportions are denoted by *X* and 1 – *X*, respectively, stationarity exists if *λ*_11_*X* = *λ*_21_(1 – *X*), i.e., *X* = *λ*_21_/(*λ*_11_ + *λ*_21_) (Bulmer 1991; McVean and Charlesworth 1999). At stationarity, the proportion of sites that are fixed for A_1_ is proportional to the expected time a site spend in that state, so that *X* =*λ*_11_^−1^/(*λ*_11_^−1^ + *λ*_21_^−1^) = *λ*_21_/(*λ*_11_ + *λ*_21_).

All the quantities needed for determining the value of Δ*π*_*e*_ in Equation (11c), other than the mutational parameters at the selected site, *θ*_*s*_ and *κ*, can be obtained from the integral formulae for the fixation probabilities, together with the simulation methods described in the previous section. The mutational parameters serve only to determine the magnitude of Δ*π*_*e*_, and have no influence on its sign, provided that the assumptions concerning the times to recovery after fixation and loss events and the relative values of *λ*_*i*0_ and*λ*_*i*1_ are met. The general equations above are valid regardless of the frequency of recombination; the values of the Δ*π* variables that appear in these expressions will, of course, be affected by recombination.

Figure 7 shows an example of estimates of Δ*π*_*e*_ for the case of a neutral locus that is completely linked to a selected locus with equal mutation rates in each direction between deleterious and favorable alleles (*κ* = 1), as a function of the scaled selection coefficient, *γ*. The deleterious allele (A_2_) has dominance coefficient *h* = 0.1, such that the relative fitnesses of heterozygotes and homozygotes for this allele are 1 – 0.1*s* and 1 – *s*, respectively. As expected from the previous results, the enhancement of diversity is maximal (approximately 0.008 with *N* = 500) when *γ* = 1.5. The fact that Δ*π*_*e*_ is significantly negative for *γ* = 0 suggests that, as noted above, the assumption of a long recovery period between successive losses or fixations at the selected locus results in an underestimation of Δ*π*_*e*_ for small *γ*.

**Figure 7.**
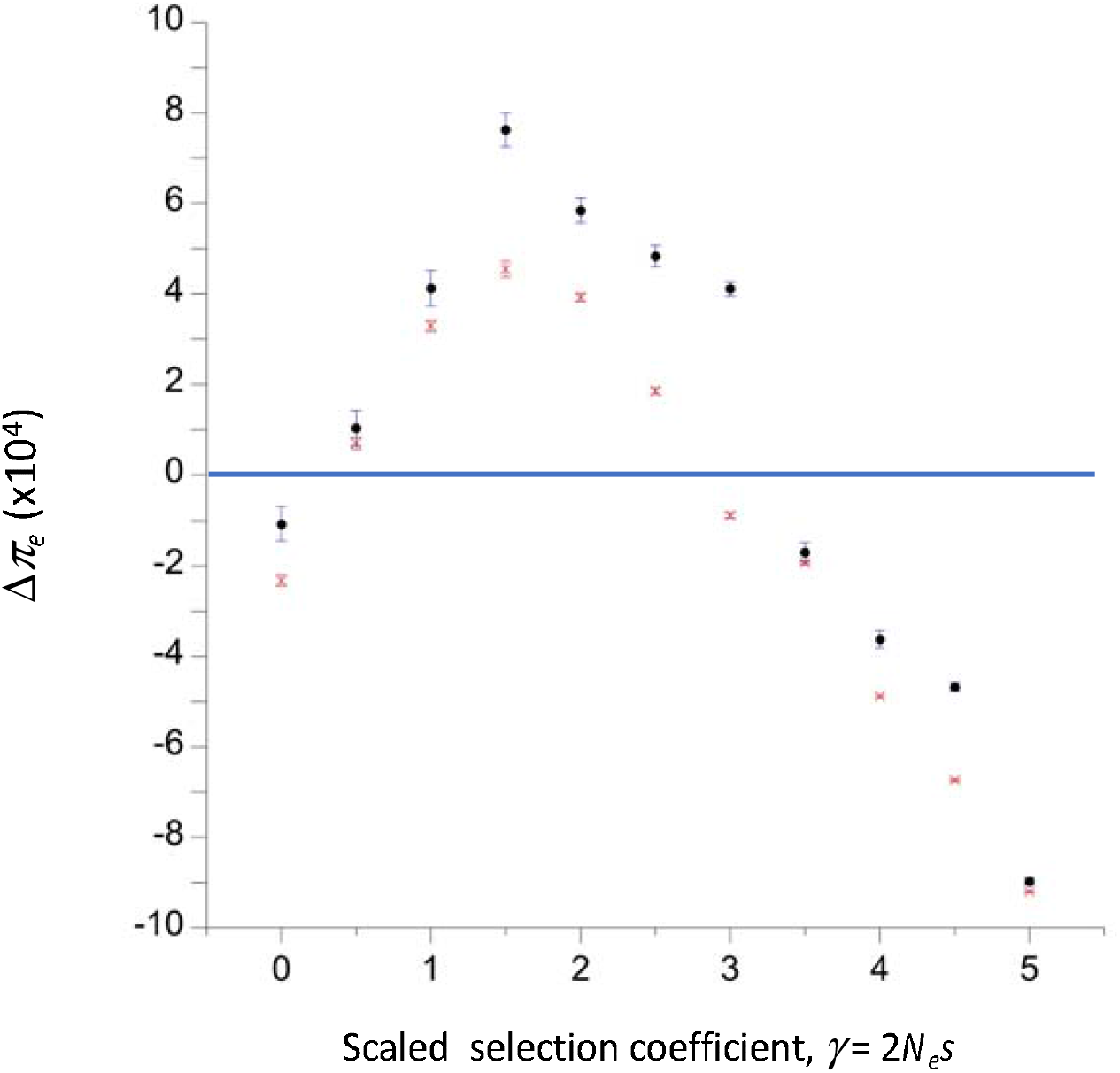
The expected deviation of relative diversity for a completely linked neutral site from its value in the absence of selection (Δ*π*_e_), at the stationary state under genetic drift, mutation and selection. The dominance coefficients for deleterious and favorable mutations are *h* = 0.1 and *h* = 0.9, respectively. Equal frequencies of mutations in each direction at the selected site were assumed, with a scaled mutation rate *θ*_s_ = 0.01. The means and standard errors over replicate simulations are shown for population sizes of 50 (red points) and 500 (black points).

In the example in the figure, the scaled mutation rate at the selected locus was 0.01 for both population sizes, corresponding to an absolute mutation rate of 5 × 10^−5^ with *N* = 50 and 5 × 10^−6^ with *N* = 500; the results are not very sensitive to the absolute value of *N* for the same set of scaled parameters, so that the approximate values for an arbitrary *θ*_*s*_ can be obtained by multiplying the results in the figure by 100*θ*_*s*_. In a population of size 10,000, the mutation rate would be 1.25 × 10^−7^, corresponding to a non-recombining region with 25 basepairs under selection with mutation rate is 0.5 × 10^−8^ per basepair, which is approximately the same as the estimate for *Drosophila melanogaster* (Assaf *et al*. 2017).

A check on the validity of this approach is provided by the case when *γ* is so large that the rate of fixation of deleterious mutations is negligible. This means that only the state when the population is fixed for the favorable allele A_1_ needs to be considered. There is then a constant input of new mutations to A_2_ alleles, which are successively lost from the population with a probability close to 1. This situation corresponds to the standard model of background selection (BGS), under which stochastic effects at the loci under selection are assumed to be negligible (Charlesworth et al. 1993). In this case, however, the assumption of a complete recovery of diversity at the neutral locus after each loss event is likely to be violated, as the large population size means that there is a high rate of input of new deleterious mutations that are then rapidly lost from the population, especially if we are dealing with a non-recombining region with numerous genes subject to purifying selection.

This problem is examined in the second part of the Appendix, where it is shown that the results obtained with the assumption of complete recovery should still provide a good approximation in this case. The deviation of mean relative diversity from one can be predicted from the sum of the Δ*π* values over the period covering the loss of an A_2_ allele and the appearance of a new mutation to A_2_, as given by Equation (10), divided by the expected time between successive losses of deleterious mutations (equivalent to multiplication by *λ*_10_), as can be seen from Equation (A12). This prediction can be compared to the standard BGS formula for autosomal loci in a randomly mating population, which assumes that allele frequencies subject to selection are unaffected by drift, i.e., *γ* >> 1. In this case, the reduction in diversity relative to neutral expectation at a neutral site completely linked to a selected locus or group of loci is approximately equal to *u*/*hs* provided that *u*/*hs* << 1, where *u* is the net rate of mutation to deleterious alleles (Charlesworth *et al*. 1993). This can be equated to Δ*π*_*e*_ for the same values of *u, h* and *s* when applying Equation (11c) to the simulations used here. To model large *γ* values while retaining the assumption of small *s*, it is necessary to use simulations with a large population size (*N* = 500). To generate these numbers, the scaled rate of mutation to deleterious mutations, *θ*_*s*_, was arbitrarily set to 1; the values for an arbitrary deleterious mutation rate of *u* can be found by multiplying these values by 4*Nu*. Some examples are shown in Table 2; it can be seen that the predictions from the present approach converge on the values from the large population size formula as *hγ* increases.

**Table 2.**
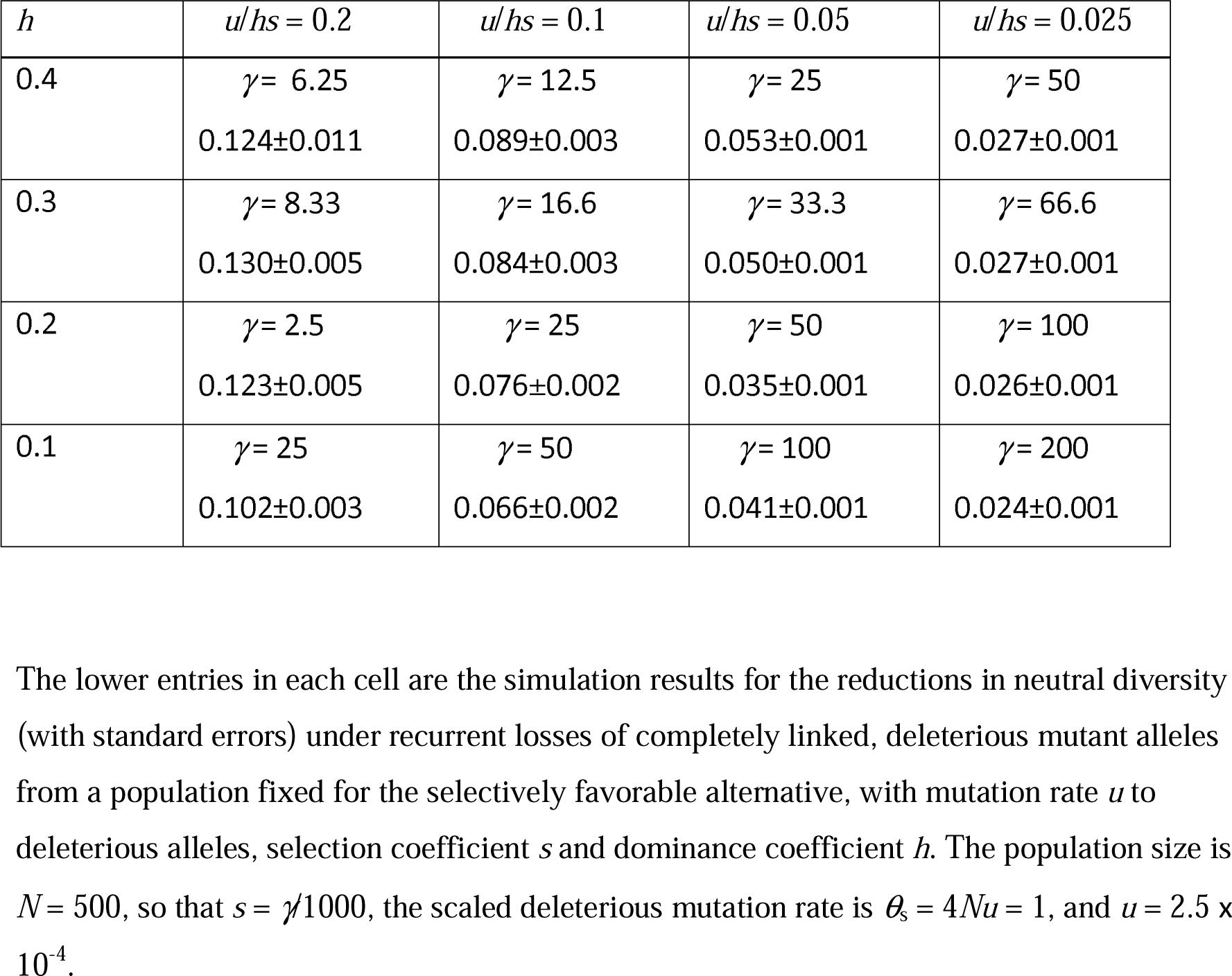
Comparison of the deterministic BGS predictions for the reduction in neutral diversity with simulations of losses of deleterious mutations from populations fixed for the favorable allele.

| $h$ | $u/hs = 0.2$ | $u/hs = 0.1$ | $u/hs = 0.05$ | $u/hs = 0.025$ |
| --- | --- | --- | --- | --- |
| 0.4 | $\gamma = 6.25$<br>0.124±0.011 | $\gamma = 12.5$<br>0.089±0.003 | $\gamma = 25$<br>0.053±0.001 | $\gamma = 50$<br>0.027±0.001 |
| 0.3 | $\gamma = 8.33$<br>0.130±0.005 | $\gamma = 16.6$<br>0.084±0.003 | $\gamma = 33.3$<br>0.050±0.001 | $\gamma = 66.6$<br>0.027±0.001 |
| 0.2 | $\gamma = 2.5$<br>0.123±0.005 | $\gamma = 25$<br>0.076±0.002 | $\gamma = 50$<br>0.035±0.001 | $\gamma = 100$<br>0.026±0.001 |
| 0.1 | $\gamma = 25$<br>0.102±0.003 | $\gamma = 50$<br>0.066±0.002 | $\gamma = 100$<br>0.041±0.001 | $\gamma = 200$<br>0.024±0.001 |

The lower entries in each cell are the simulation results for the reductions in neutral diversity (with standard errors) under recurrent losses of completely linked, deleterious mutant alleles from a population fixed for the selectively favorable alternative, with mutation rate $u$ to deleterious alleles, selection coefficient $s$ and dominance coefficient $h$ . The population size is $N = 500$ , so that $s = \gamma/1000$ , the scaled deleterious mutation rate is $\theta_s = 4Nu = 1$ , and $u = 2.5 \times$ $10^{-4}$ .

Stochastic fluctuations in the frequencies of alleles at selected loci are known to reduce the effect of BGS on diversity (Charlesworth *et al*. 1993). Table 1 of Charlesworth *et al*. (1993) gives simulation results for a range of *γ* values for the case of a non-recombining region with a deleterious mutation rate of *u* = 0.005, *s* = 0.1 and *h* = 0.2, so that *u*/*hs* = 0.25 The ratio of the negative of the natural logarithm of a simulated value of *π*/*θ* to 0.25 provides a measure of the extent of the deviation from the deterministic prediction. These measures (with their standard errors) were as follows: 0.696±0.024 (*γ* = 20), 0.892±0.035 (*γ* = 40), 0.828±0.054 (*γ* = 80) and 1.056±0.049 (*γ* = 180). The corresponding ratios from the present method for these parameter sets were 0.510±0.015, 0.858±0.020, 0.992±0.020 and 1.124±0.016, respectively. In view of the approximations needed to obtain these results, there is reasonably good agreement with the exact simulation results.

The argument concerning the effect of the times between events presented in the Appendix can be extended to the case when there is a continual flux between A_1_ and A_2_ alleles, as in the case of the model of AOD discussed above. Here, the probabilities of fixation of both variants are of the order of 1/*N*_*H*_, whereas the probabilities of loss are of the order of one. The expected numbers of loss events between successive fixations are thus of order *N*_*H*_ for both types of event. Provided that *N*_*H*_ is not too small, the argument leading to Equation (A.12) implies that the net effect on diversity is similar to the value when there is a complete recovery between loss events, so that Equation (11c) should provide a good approximation to the effects of successive losses, but there will still be an inaccuracy associated with the recovery period following a fixation event.

## Discussion

The analytical and simulation results described here shed light on the effects of the fixations and losses of weakly selected or neutral mutations on levels of genetic diversity at linked neutral sites, first studied by Tajima (1990), and which have recently received renewed attention in the population genetics literature (Mafessoni and Lachmann 2017; Johri et al. 2021; Kaushik and Jain 2021; Moinet et al. 2021). As described in the previous two sections, these effects can largely be understood in terms of the effects of the selection parameters on the times to fixation or loss, conditioned on fixation or loss, with some slight complications.

### Conditional fixation and loss times with weak selection

One might expect selection on favorable mutations to reduce their mean time to fixation relative to neutrality, and selection against deleterious mutations to have the opposite effect, with the converse applying to losses of mutations. As described above, this is not necessarily the case. Mafessoni and Lachmann (2015) used numerical integration of the relevant diffusion equation formulae for autosomal mutations with random mating to show that the mean fixation times (conditioned on fixation) of weakly selected dominant or partially dominant (*h* > ½) mutations, as well as of weakly selected, recessive or partially recessive (*h* < ½) deleterious mutations, can be larger than the corresponding neutral values. They also showed that, under similar conditions, the mean conditional times to loss of new mutations are larger than the neutral values. For sufficiently large values of the magnitude of the scaled selection coefficient, |*γ*| = 2*N*_*e*_|*s*|, however, fixation and loss times are always lower than the neutral values when there is directional selection (0 ≤ *h* ≤ 1). The approximations derived here (Equations 2 and 3), which are accurate with respect to second-order terms in *γ*, confirm these findings (see also Kaushik and Jain 2021, Equation 9). They provide a reasonably good fit to the numerical integration and simulation results for |*γ*| values of the order of 2 or less, provided that the population size is sufficiently large (Figures 1 and 2).

### Effects of weak selection on neutral diversity at linked sites

Mafessoni and Lachmann (2015) also showed that fixations of partially recessive favorable or partially dominant deleterious mutations can increase the mean level of diversity (*π*) at a linked neutral site, compared with the corresponding value for fixations of neutral mutations, if this is measured at the time of fixation. The conditions for such an increase are similar to those for an increased time to fixation (see also Johri *et al*. [2021] and Kaushik and Jain [2021]). The simulation results described here confirm this finding, and give similar results for losses of deleterious mutations (Figures 5, S4 and S5). However, the diversity levels at the times of fixation or loss are always lower than the mean equilibrium neutral diversity, *θ* = 4***N***_*e*_*µ*, even if these events involve purely neutral mutations, as was first described by Tajima (1990). As was explained in the section *Effects of weak selection on variability at linked neutral sites*, this effect is a consequence of conditioning on an unusually short coalescent time at the neutral site linked to the site under observation, and would not be expected to occur if we observe the mean diversity at this site over a long time period that includes multiple fixation or loss events. The question of whether diversity can be increased above *θ* by weak selection under appropriate conditions is thus not fully answered by these results.

A simple approximate method for evaluating the effects of successive fixations or losses at a site linked to focal neutral site was described above (Equation 10). This combines the sum of the diversities at the focal site over the generations while a fixation or loss event is in progress with the sum of the diversities over the generations that intervene until the next such event, during which diversity recovers towards the neutral equilibrium value. As shown in Figures 5, S4 and S5, the mean diversity taken over all generations during a fixation or loss event can be increased over the neutral value when |*γ*| is sufficiently small, and *h* < ½ for deleterious mutations or > ½ for favorable ones, partly due to mutational divergence between the haplotypes carrying the two different alleles at the site under selection. Under suitable conditions (a sufficiently long duration of the fixation or loss event), this effect can outweigh the reduction in diversity at the end of the fixation or loss event, resulting in a net increase in mean neutral diversity when taken over the entire period.

These considerations yield the seemingly paradoxical result that neutral diversity can be enhanced by fixations or losses of weakly selected favorable or deleterious mutations, in contrast to the usual assumption that diversity is reduced by fixations of favorable mutations (selective sweeps) or losses of deleterious mutations (background selection). It has, however, long been known that neutral diversity can in principle be increased by selection against deleterious recessive or partially recessive mutations at closely linked sites – the process of associative overdominance or AOD (Frydenberg 1963; Ohta 1971; Latter 1998; Pamilo and Palsson 1998; Palsson and Pamilo 1999; Wang and Hill 1999; Becher et al. 2020; Gilbert et al. 2020; Waller 2021). A correct analytical treatment of this process for the simplest case of a single selected locus and a linked neutral locus has, however, only recently been provided (Zhao and Charlesworth 2016). As shown in Figure 4, the critical dominance coefficient for the operation of AOD versus background selection (BGS) as a function of |*γ*| has similar, but not identical, properties to the conditions for increases over the neutral value of the mean conditional fixation and loss times of new deleterious or favorable mutations.

The section *Relevance to associative overdominance and background selection* describes how to relate the simulation results for the effects of selection on the diversity statistics at a linked neutral locus, when there is a continual flux of mutations between sites fixed for deleterious and favorable mutations. Under this scenario, the net mean deviation from one of *π*/*θ* is given by Δ*π*_*e*_ in Equation (11c). Fixations and losses contribute approximately equal amounts to Δ*π*_*e*_; although the effects on diversity of losses are much smaller than those of fixations, this is offset by the much higher frequencies of loss events.

The results described here provide a new perspective on the relation between AOD and BGS. If one imagines a locus subject to reversible mutation between a wild-type allele (A_1_) and a deleterious allele (A_2_), the standard model of BGS corresponds to the situation when selection against A_2_ is so strong (|*γ*| >>1) that there is effectively no possibility of fixations of reverse mutations to A_1_, and the system can be treated as though it is at equilibrium under mutation and selection, as in the standard formulae used to describe the effect of BGS on neutral diversity (Charlesworth, et al. 1993; Hudson and Kaplan 1995; Nordborg, et al. 1996). In reality, the population size is finite, so mutations from A_1_ to A_2_ are constantly occurring and being quickly lost from the population, with a probability close to one when |*γ*| >>1. As shown in Table 2, the application of the approach described here to this situation closely approximates the properties of BGS when finite population size is modeled by exact simulations.

If |*γ*| is sufficiently small, however, the effects of genetic drift and reversible mutation between A_1_ and A_2_ need to be considered jointly, as in the Li-Bulmer model of selection on codon usage (Li 1987; Bulmer 1991; McVean and Charlesworth 1999). With |*γ*| I≤ 2 or so and *h* for deleterious mutations less than ½, the results enshrined in Equations (11) show that diversity at linked neutral sites can be increased over neutral expectation when the population is at statistical equilibrium under drift, mutation and selection for sites under selection. These findings complement the results of the analysis of AOD by Zhao and Charlesworth (2016), who examined the rate of loss of neutral variability in a population that was initially at statistical equilibrium, and then placed into an environment with a relatively small ***N***_*e*_. As noted already, the conditions on the dominance coefficient as a function of |*γ*| that allow an enhancement of variability are similar for the two approaches. The framework described here is more appropriate than that of Zhao and Charlesworth (2016) for populations maintained for a long time at low effective population sizes, as is the case in some laboratory experiments that have yielded evidence for AOD (Latter 1998), and for low recombining genomic regions in large populations, where recent work has also suggested the operation of AOD (Becher *et al*. 2020; Gilbert *et al*. 2020).

### Interpreting the effects of selection on the conditional sojourn times, *t*\* and *t*\*\*

Here, intuitive interpretations are presented of the conditions for increases in the mean conditional times to fixation (*t*\*) and loss (*t*\*\*) over their neutral expectation. A more rigorous approach to this question is given in Supplementary File S1, section 3. One important point to note is that Equations (2) and (3) show that the first-order terms in *γa* in the expressions for *t*\* and *t*\*\* are zero when *b* = 0 (*h* = ½ for autosomal mutations), so that the conditional sojourn times are then approximately the same as with neutrality when selection is very weak. This finding suggests that the first-order effects of selection that depend on the term in *γa* in Equation (1) are absorbed into the probabilities of fixation and loss, so that the conditional sojourn times are controlled by *γb* alone when selection is very weak. Another relevant point is that the relative effectiveness of selection versus drift can be quantified by dividing Equation (1) by *x*(1 – *x*), which gives the ratio of the selective change in allele frequency per generation to the sampling variance under drift, *x*(1 – *x*)/2*N*_*e*_. This ratio is denoted here by *f*(*x*) = *γ* (*a* + *bx*). If *γb* < 0, the relative effectiveness of selection in causing a movement of *x* away from zero, as measured by *f*(*x*), is reduced, compared with the quasi-neutral case with *γb* = 0, and decreases as *x* increases.

The condition for weak selection to cause an increase in *t*\* over neutrality is then relatively easy to interpret (see Mafessoni and Lachmann (2015) for a different viewpoint). If *γ* > 0 and *b* < 0 (*h* > ½ for an autosomal mutation), *x* tends to be decreased by selection compared with quasi-neutrality, especially at larger values of *x*, so that the net time to reach fixation is increased by selection, and the times spent at larger values of *x* are increased relative to the neutral case. If *γ* < 0 and *b* > 0 (*h* < ½ for an autosomal mutation), there is a stronger tendency for *x* to move towards zero than under quasi-neutrality; because we are conditioning on fixation of A_2_, this causes the sojourn time to be increased over neutrality. The pull towards lower values of *x* is weaker for small *x*, so that the time spent at low values of *x* is increased relative to the neutral case. These effects on the sojourn time densities at different frequencies are illustrated in the lower panels of Figure 8.

**Figure 8.**
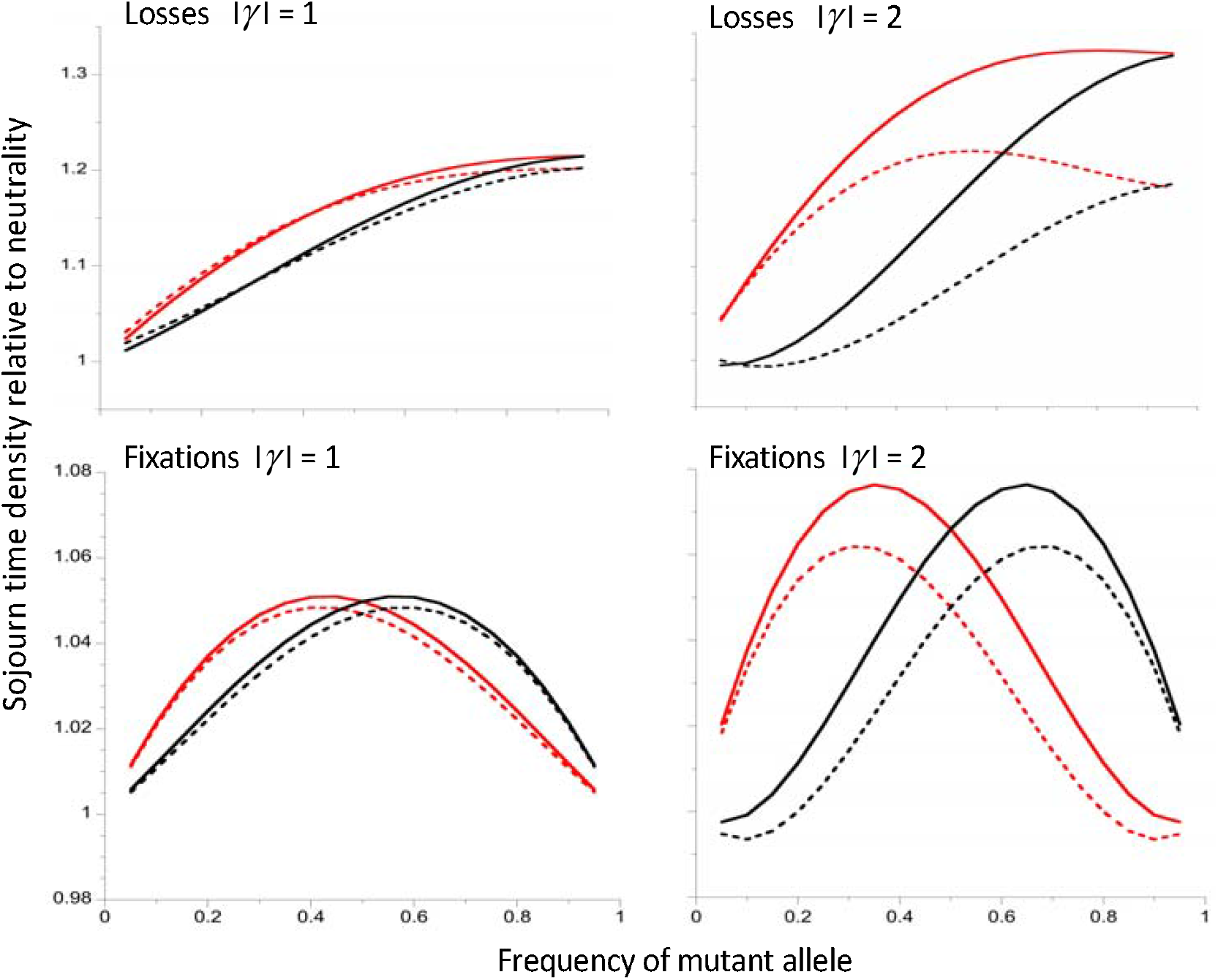
The upper two panels show the approximate (full curves) and exact (dashed curves) sojourn time densities for mutations that become lost, relative to the corresponding neutral values. The lower two panels are the sojourn time densities for mutations that become fixed. The left-hand panels are for |*γ*| = 1 and the right-hand panels are for |*γ*| = 2. Deleterious mutations with *h* = 0.1 are the red curves and beneficial mutations with *h* = 0.9 are the black curves. The exact results were obtained by numerical integrations of the relevant equations described in the Appendix; the approximate results were obtained from Equations (A5) and (A7).

It is less easy to interpret the properties of *t*\*\*, because the argument used for *t*\* would at first sight suggest a reduction rather than an increase in *t*\*\* when *γb* < 0. The most plausible interpretation is that losses of new mutations are largely caused by drift, and take place rapidly as long as *x* remains close to zero. However, the time for an A_2_ mutation to reach a relatively high frequency is increased by weak selection when *γb* < 0. It is then relatively immune to loss, and hence spends longer segregating in the population. This interpretation is consistent with the fact that the sojourn time densities relative to the neutral values are increasing functions of *x* for both favorable and deleterious mutations, as is shown in the upper panels of Figure 8.

### Conclusions and future prospects

The work described here illuminate the conditions under which weak directional selection can interact with genetic drift to cause an increase rather than a decrease in genetic diversity at linked neutral sites. It also sheds light on the conditions under which associative overdominance rather than background selection operates. The results are, however, limited in several important respects. First, only the pairwise diversity measure *π* has been studied, so that the properties of the site frequency spectra at neutral sites linked to the target of selection have not been examined; the results of Mafessoni and Lachmann (2015) and Johri *et al*. (2021) suggest that an increase in variability at a neutral locus caused by fixations of linked weakly selected mutations is accompanied by a reduction in the frequencies of rare variants relative to the standard neutral expectation. Further work is needed to investigate the magnitude and direction of distortions of the site frequency spectrum in the context of recurrent fixations and losses of weakly selected mutations. These events can cause associative overdominance, which is expected to cause a skew toward intermediate frequency variants (Becher et al. 2020; Gilbert et al. 2020).

Second, the extent to which multiple weakly selected loci will mutually affect each other has largely been unexplored; a recent review of relevant data suggests that such effects of multiple loci may be important for explaining the unexpectedly high levels of inbreeding depression and variation in quantitative traits in small populations (Waller 2021). Previous simulation work has shown that, with low rates of recombination and many tightly linked highly recessive or deleterious mutations, the population can “crystallize” into two haplotypes carrying complementary sets of mutations that exhibit pseudo-overdominance (Charlesworth and Charlesworth 1997; Palsson 2000; Gilbert et al. 2020); there is evidence for less severe effects, causing a retardation of loss of variability, when multiple loci are subject to deleterious mutations in small populations (Latter 1998; Bersabé *et al*. 2016).

Further investigation of this topic is desirable. It would also be of interest to investigate the properties of subdivided populations, where stochastic effects on loci under selection can be significant when deme sizes are sufficiently small (Roze and Rousset 2004; Roze 2015; Charlesworth 2018).

Finally, the properties of fixation times in varying environments can be significantly different from those for a constant environment that are used here, with a breakdown in the symmetry between *h* and 1 – *h* for favorable and deleterious mutations as far as *t*\* is concerned (Kaushik and Jain 2021). It is likely that this will also apply to times to loss. The relations of these quantities with associative overdominance in small laboratory populations will probably not be affected by this effect, but the behavior of natural populations could be significantly changed.

## Data Availability

No new data or reagents were generated by this research. The codes for the computer programs used to produce the results described below will be made available on Figshare on acceptance.

## Conflicts of Interest

None

## Acknowledgments

I thank two anonymous reviewers, Graham Coop, Kavita Jain and Lei Zhao for their comments on the manuscript, which have helped to improve it; in particular, Lei Zhao detected errors in the original versions of Equations (3b) and (8b), which have been corrected.

## Appendix

### Conditional sojourn time formulae

Equation (4.52) of Ewens (2004) can be used to evaluate the sojourn time density conditional on fixation for an allele A_2_ at frequency whose initial frequency is *q* = 1/*N*_*H*_, so that *x* ≥ *q*. This equation can be rewritten as:

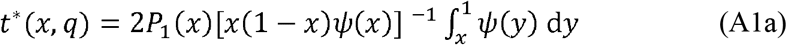

where *P*_1_(*x*) is the probability of fixation from frequency *x*, respectively, which is given by:

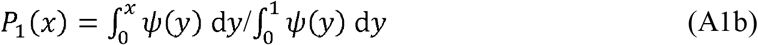

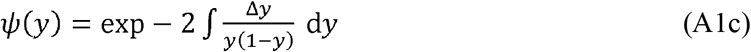

Similarly, the probability of loss from frequency *x* is given by:

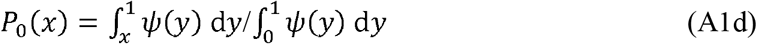

(Ewens 2004, Equations 4.15-4.17).

Substituting the expression for Δ*x* given by Equation (1) of the main text into Equation (A1c) we have:

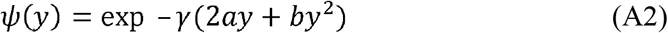

For *b* = 0 (semi-dominance) analytic expressions for the integrals in Equations (A1) are available (Ewens 2004, pp.165, 169-170). In other cases, numerical evaluations are needed if the approximations derived below are to be avoided. For this purpose, it is convenient to use series representations of the integrals in Equations (A1b) and (A1c). Expanding the exponential function in Equation (A2), the indefinite integral of *ψ*(_y_) can be written as:

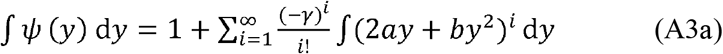

With *b* ≠ 0, and writing *c* = *a*/*b*, this expression can be reduced to a simpler form by the transform 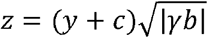, which gives 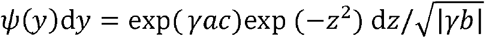 if *γb* > 0 and 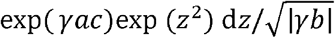 if *γb* > 0. The terms in exp (−*z*^2^) or exp (*z*^2^) can then be expanded as a Taylor series in *z*, and integrated term by term to yield an infinite series in *z*:

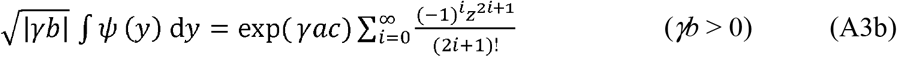

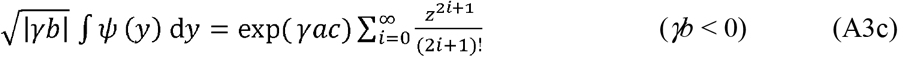

If third- and higher-order terms in *γ* in Equations (A1) - (A3a) are neglected, the following approximations for the components of Equations (A1) emerge after some algebra:

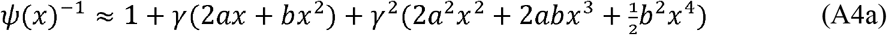

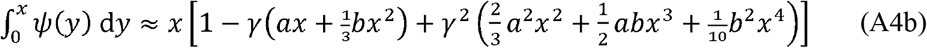

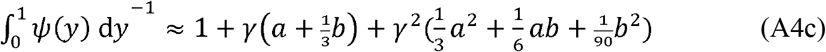

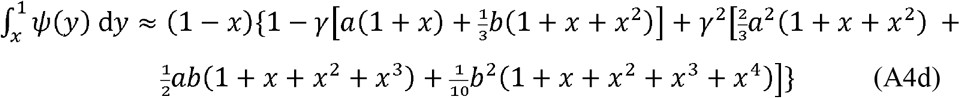

Inserting these expressions into Equation (A1a), and neglecting higher-order terms in *γ*, we obtain:

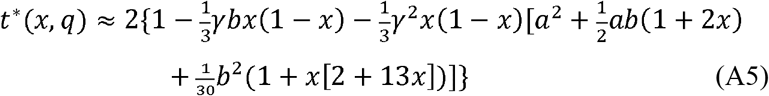

For the sojourn time density conditional on loss, Equation (4.52) of Ewens (2004) can be used:

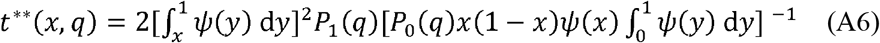

Using the procedure applied to *t*\*(*x, q*), the following second-order approximation with respect to *γ* is obtained:

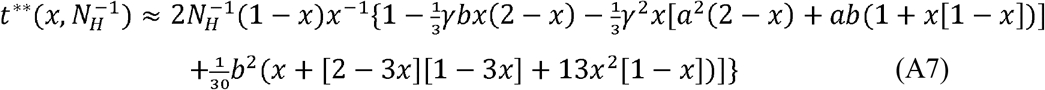

An approximation for the density function for the expected sojourn time between loss or fixation at frequency *x* for a new mutation can similarly be obtained from Equation (4.23) of Ewens 2004):

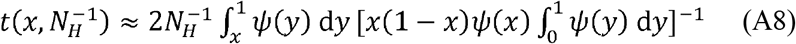

This expression yields:

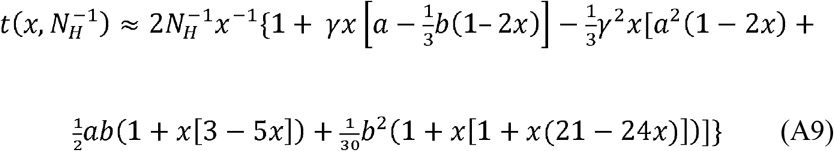

### Background selection approximation

In this case, only losses of new mutations to the deleterious allele A_2_ from a population fixed for the favorable allele A_1_ need to be considered. It is assumed that this process has been going on indefinitely, and that the population is being observed over a long time period *T*_*o*_ (in units of 2*N*_*e*_ generations); the mean diversity at the neutral site is then taken over this period. From Table 1, the expected number of losses of A_2_ over this period is *λ*_10_*T*_*o*_ = 0.5*θ*_*s*_*N*_*H*_*P*_10_*T*_*o*_, and the expected time interval between successive losses is *T*_*l*_ = 1/*λ*_10_, where *T*_*l*_ << 1 in the case of BGS. At the start of this process, which can be assumed to have occurred long before the period of observation, there is a deviation Δ*π*_2*l*_ from 1 of the relative diversity at the end of the first loss event (see Table 1), whose value can be found by the simulation procedure described in the main text. This is followed by a recovery period, whose length is drawn from an exponential distribution with rate parameter *λ*_10_, so that the new expected value of Δ*π* at the end of the recovery period is:

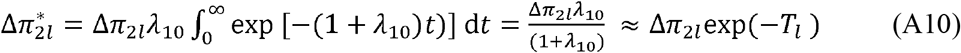

The sum of the Δ*π* values over the recovery period is approximately equal to Δ*π*_2*l*_*T*_*l*_, since most values will be close to Δ*π*_2*l*_ due to the short length of this period; a more exact derivation of this result can be obtained by use of the argument that yielded Equation S22 of Campos and Charlesworth (2019).

If the change in diversity at each event is small, we can assume additivity of individual effects in order to obtain the net change after several events. The second loss event thus results in a deviation before recovery of 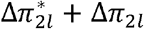 and a corresponding sum of Δ*π* over the recovery period of 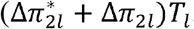. Using the argument above, the post-recovery deviation ≈ 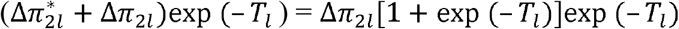, and the corresponding sum of Δ*π* values over the recovery period ≈ Δπ_2*l*_ (1+ exp(– *T*_*l*_)]*T*_*l*_. If this process is repeated indefinitely, it can be seen that the individual post-recovery deviations are given by the product of Δπ_2*l*_ exp (– *T*_*l*_) and the sum of a geometric series in exp (− *T*_*l*_), which converges on:

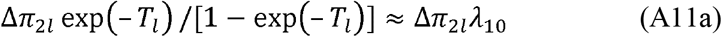

Similarly, the sum of the individual deviations over the recovery period converges on:

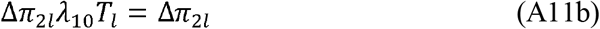

This formula is the same as for the case when there is a complete recovery of diversity after a loss of A_2_. It therefore follows that the net deviation in diversity caused by multiple losses of A_2_ at rate *λ*_10_ is approximated by:

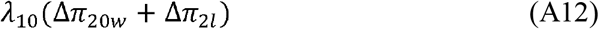

## Notes

### Competing Interest Statement

The authors have declared no competing interest.

### Summary of Updates

Some of the less important details have been removed to a supplementary file, and a number of small errors corrected.

